# Multi-scale photocatalytic proximity labeling reveals cell surface neighbors on and between cells

**DOI:** 10.1101/2023.10.28.564055

**Authors:** Zhi Lin, Kaitlin Schaefer, Irene Lui, Zi Yao, Andrea Fossati, Danielle L. Swaney, Ajikarunia Palar, Andrej Sali, James A. Wells

## Abstract

The cell membrane proteome is the primary biohub for cell communication, yet we are only beginning to understand the dynamic protein neighborhoods that form on the cell surface and between cells. Proximity labeling proteomics (PLP) strategies using chemically reactive probes are powerful approaches to yield snapshots of protein neighborhoods but are currently limited to one single resolution based on the probe labeling radius. Here, we describe a multi-scale PLP method with tunable resolution using a commercially available histological dye, Eosin Y, which upon visible light illumination, activates three different photo-probes with labeling radii ranging from ∼100 to 3000 Å. We applied this platform to profile neighborhoods of the oncogenic epidermal growth factor receptor (EGFR) and orthogonally validated >20 neighbors using immuno-assays and AlphaFold-Multimer prediction that generated plausible binary interaction models. We further profiled the protein neighborhoods of cell-cell synapses induced by bi-specific T-cell engagers (BiTEs) and chimeric antigen receptor (CAR)T cells at longer length scales. This integrated multi-scale PLP platform maps local and distal protein networks on cell surfaces and between cells. We believe this information will aid in the systematic construction of the cell surface interactome and reveal new opportunities for immunotherapeutics.

## Introduction

Cell surface proteins are critical mediators of information, nutrients, and functions on cells and between them. The extracellular proteome, both secreted and membrane-bound, is encoded by more than 25% of the human genome (*1, 2*). Proteomics methods have made great strides in characterizing the composition of the surface proteome in health and disease models (*3, 4*). However, we know much less about the protein-protein interactions formed on the cell membrane, especially transient interactions that regulate cell signaling networks.

Proximity labeling proteomics (PLP) methods have enabled the identification of protein interactomes in complex cellular environments (*5, 6*). These methods typically generate a single reactive intermediate locally to label and profile nearby proteins using imaging or proteomics. The first generation of PLP methods used genetically encoded enzymes such as APEX (*7*), BioID (*8*) or TurboID (*9, 10*) to produce phenoxyl radicals or activated AMP that have long reactive half-lives (t_1/2_>100 µsec). These methods are well-suited for characterizing cell-cell and organelle-specific interactomes given their long labeling range up to 3000 Å by labeling electron-rich amino acids (*11*). Singlet oxygen generators (SOG) triggers selective labeling on His (*12*) at a shorter range given the shorter half-life of singlet oxygen in water (∼2-4 μs) (*13*). However, proteins are estimated to be separated by only 60-70 Å on the crowded cell surface (*14*), thus making it challenging to identify the most proximal protein neighbors using long-range PLP approaches.

Most recently, PLP methods of very short range have emerged, enabling higher resolution mapping including µMap (*15–17*). These designs employ transition-metal or other photocatalysts attached to antibodies to trigger reactive intermediate with shorter half-lives such as carbenes or nitrenes (t_1/2_ ∼2 and 10 ns, respectively) (*11, 18*). Activation of these probes enables labeling proteins at a significantly shorter range of ∼100-700 Å as well as broader amino acid coverage (*11, 19*), thus making it much more appropriate for nearest neighborhood analysis (*20–23*). Collectively, the suite of PLP methods can cover a broad length scale for labeling protein neighborhoods and synapses but require multiple photocatalysts for adjustable resolution.

Here, we report a multi-scale photocatalytic PLP technology, termed MultiMap (**Fig. 1A**) that allows short-, intermediate-, and long-range labeling from a single photocatalyst, Eosin Y (EY). We discovered that EY, a fluorescent dye commonly used in food chemistry and biological staining (*21*) can efficiently trigger the generation of carbene, nitrene, singlet oxygen and phenoxyl radicals from photo-probes with bio-compatible blue or green light. We applied MultiMap to profile high-resolution neighborhoods of the oncogenic epidermal growth factor receptor (EGFR) in different cellular contexts. We identified >20 neighbors and further validated their interactions via immunoprecipitation and *in silico* prediction models using AlphaFold-Multimer (*24*). We demonstrated that MultiMap can capture long-range intercellular engagements between cancer cells and T lymphocytes induced by bi-specific T-cell engagers (BiTEs) and engineered chimeric antigen receptors (CARs). Our data show that MultiMap is an effective multi-scale PLP technology that can characterize local and distal cellular interactomes from a single photocatalyst. We believe that with artificial intelligence-assisted structural prediction methods integrated, the MultiMap workflow will be an important approach in the broad quest to define the spatial organization of the cell surface proteome and to reveal new drug discovery opportunities.

**Fig. 1.**
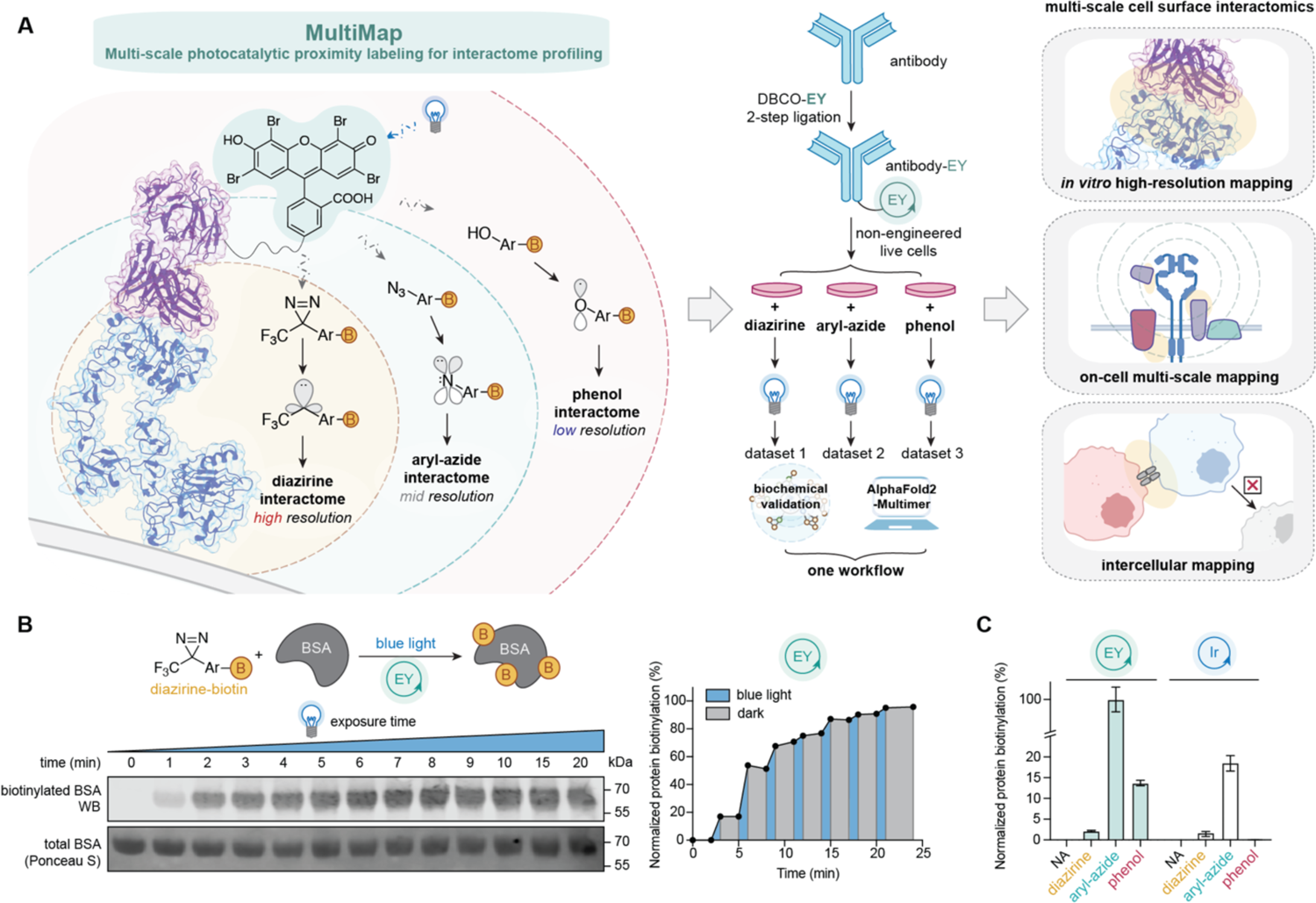
MultiMap captures high-resolution snapshots of biological networks across a wide range of length scales. **(A)** A schematic of the MultiMap workflow. EY is conjugated to an antibody that binds the target of interest (e.g., Fab arm of Ctx bound to the EGFR extracellular domain). Upon illumination, proteins are biotinylated, captured and digested for MS analysis. Proteomics hits are further examined by immunoprecipitation and predictive structural analysis via AlphaFold-Multimer. MultiMap is a useful platform for profiling local membrane protein interactomes both on live cells and between cell-cell synapses. **(B)** WB showing EY-mediated photocatalytic biotinylation of BSA by a diazirine-biotin probe upon blue LED illumination. Biotinylation can be controlled temporally by pulsed light. **(C)** EY triggers labeling of BSA with all three photo-probes (diazirine-biotin, aryl-azide-biotin and phenol-biotin), but Ir only activates the first two. All immunoblot images are representative of at least two biological replicates.

## Results

### Eosin Y (EY) activates a panel of photo-probes for protein labeling

We explored Eosin Y (EY) for PLP (**Fig. 1A**) based on its photocatalytic ability broadly used in polymer synthesis (*25*) and easy commercial access (*26*). In contrast, the current transition-metal photocatalysts used in µMap, such as {Ir[dF(CF_3_)ppy]_2_(dtbbpy)}PF_6,_ requires lengthy synthetic routes (*15, 27*). We first compared EY to the Ir-catalyst for photo-induced hydrolysis of the diazirine via LC-MS and observed 100% quantitative hydrolysis with 10 min blue LED illumination (λ=450 nm, **fig. S1A**). The absorption peak for EY (λ_max_=517 nm, **fig. S1B**) is significantly red-shifted compared with the Ir-catalyst (λ_max_=420 nm) (*15*), which could make EY more bio-compatible (*28*). Indeed, green LED illumination (λ=525 nm) of EY induced 100% diazirine hydrolysis, whereas the iridium catalyst did not induce any detectable conversion (**fig. S1A**). Erythrosin B (λ_max_=535 nm), a dye structurally similar to EY, photo-catalyzed 31% hydrolysis of the diazirine with blue LED illumination and 100% with green LED illumination, indicating that the EY scaffold can be modified for specific photochemical properties. We next tested the ability of EY to label bovine serum albumin (BSA) using a diazirine-biotin probe in the presence of light (**Fig. 1B**). We observed time- and light-dependent accumulation of biotinylated BSA via Western blot (WB) analysis; labeling plateaued within 6 min of blue LED illumination (**Fig. 1B** left). A pulse-light experiment (**Fig. 1B** right) demonstrated that the catalytic function of EY is light-dependent.

EY is structurally similar to other photocatalysts that can trigger aryl-azide and phenol probes (*26, 29, 30*). EY fully converted the aryl-azide probe to the aniline upon blue LED illumination (**fig. S1C**). We then tested the photocatalytic labeling on BSA by WB analysis and found that EY efficiently catalyzed biotin labeling in the presence of either aryl-azide-biotin or phenol-biotin (**Fig. 1C** and **fig. S1D-F**). Additionally, we evaluated the ability of EY to induce singlet-oxygen-based labeling using biocytin-hydrazide (*12*) and observed efficient light-dependent labeling. The extents of labeling of BSA among the four biotin-containing reactive probes, which we refer to as photo-probes, ranged in the following order: aryl-azide-biotin (>95%), biocytin-hydrazide (∼80%), phenol-biotin (∼15%) and diazirine-biotin (∼2%) (**fig. S1D-E**). Under blue light illumination, the Ir catalyst could trigger labeling of BSA by the aryl-azide-biotin but not the phenol-biotin (**fig. S1D-E**). EY also efficiently catalyzed labeling of BSA with green LED while the Ir catalyst showed no labeling (**fig. S1F**). More than 80% labeling of BSA was achieved upon 3 min green LED exposure of EY with all four photo-probes. We also found that EY maintains its photocatalytic function above its pKa (pH=3.5) (**fig. S1G-H**) (*26*) and thus is compatible with labeling across a wide range of physiologic pH conditions.

### Conjugation of EY onto proteins

We evaluated different conjugation methods for EY first onto BSA and then to antibodies (**Fig. 2A** and **fig. S2A**). After synthesizing DBCO-PEG_4_-EY via an amine-isothiocyanate reaction (**Scheme 1**), we explored the conjugation efficiency and stoichiometry for attaching a click-compatible azido functionality specifically to Lys, Met or Cys using N-hydroxy succinimide (NHS) ester, oxaziridine, or maleimide/iodoacetamide warheads, respectively (**fig. S2B**). EY-conjugation via NHS-azide ligation produced the most efficient conjugation; conjugated EY also efficiently catalyzed BSA self-biotinylation with diazirine-biotin, aryl-azide-biotin and phenol-biotin (**fig. S2B-C**).

**Fig. 2.**
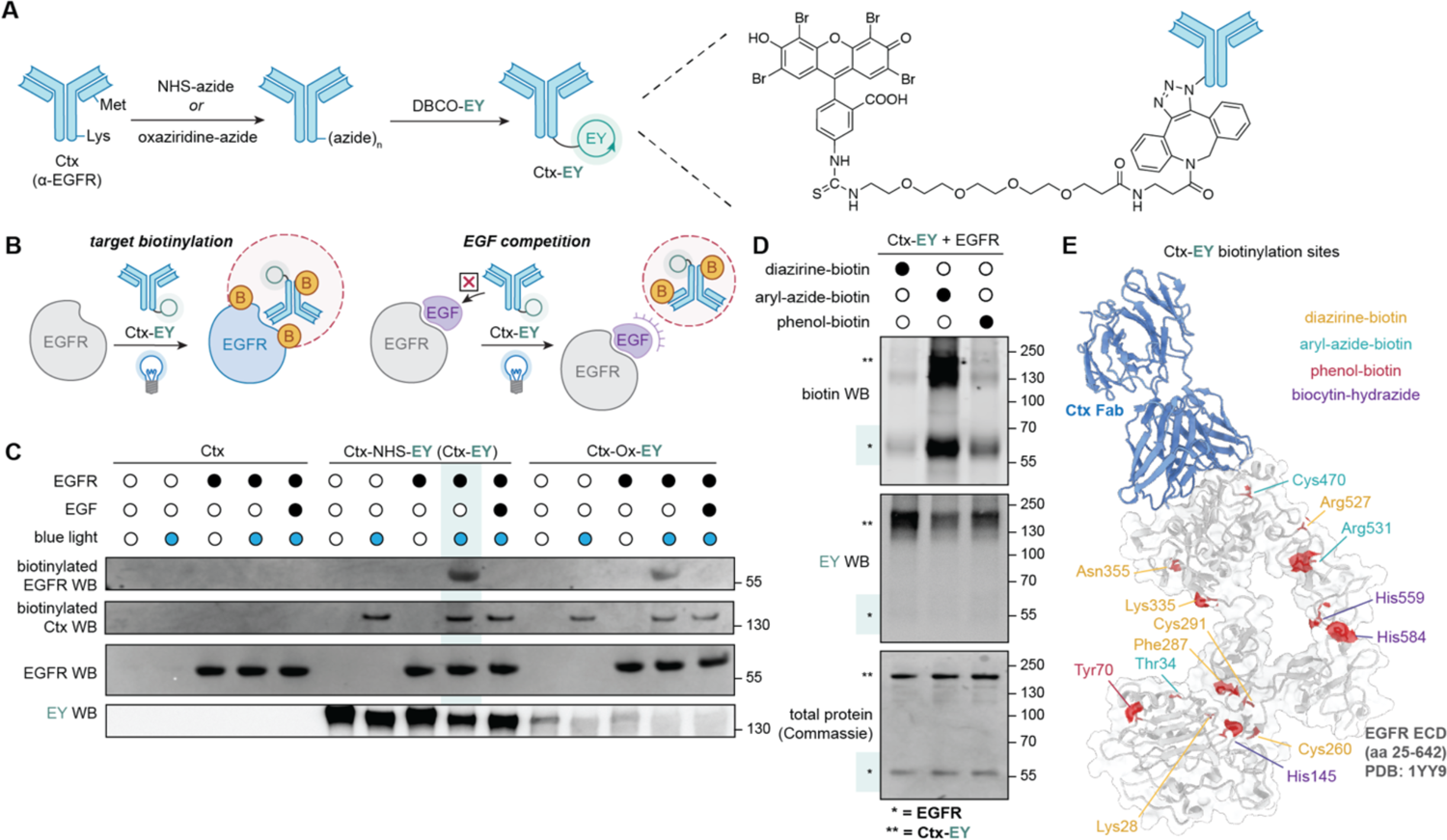
Targeted labeling using antibody-EY conjugates *in vitro*. **(A)** Synthetic scheme of Ctx-EY via a two-step bioconjugation workflow. An azido functionality was first introduced onto either Lys or Met residues using NHS or oxaziridine chemistry, respectively, followed by bio-orthogonal click reaction to couple EY. **(B)** Schematic design to test intra- and inter-biotinylation of Ctx-EY and EGFR with or without EGF competition. **(C)** Targeted EGFR biotinylation with the diazirine-biotin photo-probe when triggered by either Ctx-NHS-EY (Ctx-EY) or Ctx-Ox-EY *in vitro*. Both conjugates selectively label EGFR in a light-dependent fashion, which is competed off by exogenous EGF. **(D)** EGFR is biotinylated by all three photo-probes: diazirine-biotin, aryl-azide-biotin or phenol-biotin using Ctx-EY. **(E)** Biotinylation sites of diazirine-biotin (yellow), aryl-azide-biotin (cyan), phenol-biotin (maroon) and biocytin-hydrazide (purple) highlighted on the crystal structure of the EGFR ECD (grey) in complex with Ctx Fab (blue) (PDB: 1YY9, data taken from **Table S5-8**). All immunoblot images are representative of at least two biological replicates.

Next, we conjugated EY to cetuximab (Ctx), an FDA-approved antibody that selectively binds EGFR and competes for epidermal growth factor (EGF) binding, thus turning-off EGFR signaling and cell proliferation in cancer (**Fig. 2B**) (*31, 32*). Ctx does not have Lys, Met or Cys residues in the CDRs or in the contact epitope with the EGFR ectodomain (ECD, aa 1-645, PDB:1YY9) (*31*), suggesting all bioconjugation methods are viable without impairing binding. We tested the same panel of bioconjugation warheads on Ctx, generating similar levels of conjugation as seen for BSA (**fig. S2D**). Quantification of the levels of conjugation by WB analysis or EY absorption indicated that a stoichiometry of eight and two EY catalysts were installed per Ctx-NHS-EY and Ctx-Ox-EY, respectively (**fig. S2E-F**).

We then tested the intra- and inter-molecular labeling of the Ctx-EY conjugates with equimolar amounts of recombinant human EGFR ectodomain (ECD, aa 1-645) and in competition with EGF (**Fig. 2C**). 1 µM Ctx-NHS-EY or Ctx-Ox-EY was mixed with 1 µM EGFR-ECD with or without pre-incubation with 1 µM EGF. 100 µM diazirine-biotin was added and the mixture was illuminated with blue LED for 10 minutes. Proteins were purified via acetone precipitation and immunoblotted to evaluate the labeling efficiency (**Fig. 2C**). Both Ctx-NHS-EY and Ctx-Ox-EY conjugates demonstrated self-labeling in a light-dependent manner. Not surprisingly, there was a higher degree of biotinylation with Ctx-NHS-EY which contains ∼4-fold more conjugated EY than Ctx-Ox-EY (**fig. S2D**). Intermolecular EGFR labeling with both EY-conjugated constructs occurred in a light-dependent manner (**Fig. 2C**), indicating that the conjugation of EY did not interfere with Ctx binding to EGFR as expected. Pre-incubation of EGF prevented labeling, demonstrating that direct binding is necessary for target labeling (**Fig. 2C**). To explore the generality of the workflow, we performed the same NHS and oxaziridine bioconjugation and labeling using a trastuzumab (Trz) Fab that binds the extracellular domain of the HER2 receptor (**fig. S3A**). Similar intermolecular labeling of HER2 was observed with Trz-NHS-EY or Trz-Ox-EY in a light-dependent manner. The demonstration of EGFR and HER2 labeling *in vitro* supports the broad applicability of the bioconjugation strategy and photo-probe labeling workflow.

We chose to focus on the NHS-azide conjugate (abbreviated to Ctx-EY) given its higher bioconjugation and photo-probe labeling efficiency. We evaluated the EGFR labeling efficiencies with diazirine-, aryl-azide-, and phenol-biotin photo-probes in parallel (**Fig. 2D**). All three probes labeled the EGFR ECD to increasing levels: aryl-azide-biotin> phenol-biotin> diazirine-biotin. The differing yields likely result from the combined effects of the reactive radical intermediates: half-lives (phenol>>aryl-azide>diazirine), yield of reaction with protein (phenol>aryl-azide>diazirine), and broad amino-acid preference observed (diazirine∼aryl-azide>>phenol) (**fig. S3B**) (*33–35*).

We analyzed the specific sites of biotinylation for self-labeling of BSA and binary complex labeling of Ctx and EGFR with different photo-probes using MS analysis (**Table S1-8** and **fig. S3B-C**). For diazirine-biotin, we found 17 biotinylated sites on BSA (**Table S1**), and 30 sites on the Ctx-EGFR complex (**Table S5**), with good coverage of modified peptides over the light and heavy chains of Ctx, as well as EGFR ECD (**Fig. 2E** and **fig. S3C**). We further characterized the modification sites on BSA and Ctx-EGFR systems for the other probes (**Table S1-8** and **fig. S3B-C**). As expected, phenol-biotin mostly labeled Tyr/Trp, while labeling with biocytin-hydrazide was found exclusively on His. Diazirine-biotin and aryl-azide-biotin showed very broad amino acid preference, consistent with previous reports (*19, 34*).

### Ctx-EY catalyzes targeted labeling of EGFR on cells

We next evaluated the ability of Ctx-EY to bind EGFR and label live cells (**Fig. 3A**). First, we incubated the Ctx-EY conjugate with an epithelial skin cancer cell line, A431 cells, that endogenously expresses very high levels of wild-type EGFR (nTPM =2978) (*36*). On-cell binding for the Ctx-conjugates, both Ctx-EY and Ctx-Ir, was confirmed via flow cytometry showing that the conjugation of EY or the Ir-catalyst did not affect binding (**Fig 3B**). Detailed titration from 1 nM to 10 μM of Ctx and Ctx-EY analyzed via flow cytometry further confirmed conjugation did not detectably affect cell binding (**Fig. 3B** and **fig. S4A**). We also tested A549 cells with more typical levels of EGFR (*37*) (nTPM=59.7, **fig. S4B**) as well as NCI-H441 cells with very low EGFR expression (nTPM=29.8, **fig. S4C**), both of which showed proportionally reduced binding of Ctx-EY and was similar to Ctx.

**Fig. 3.**
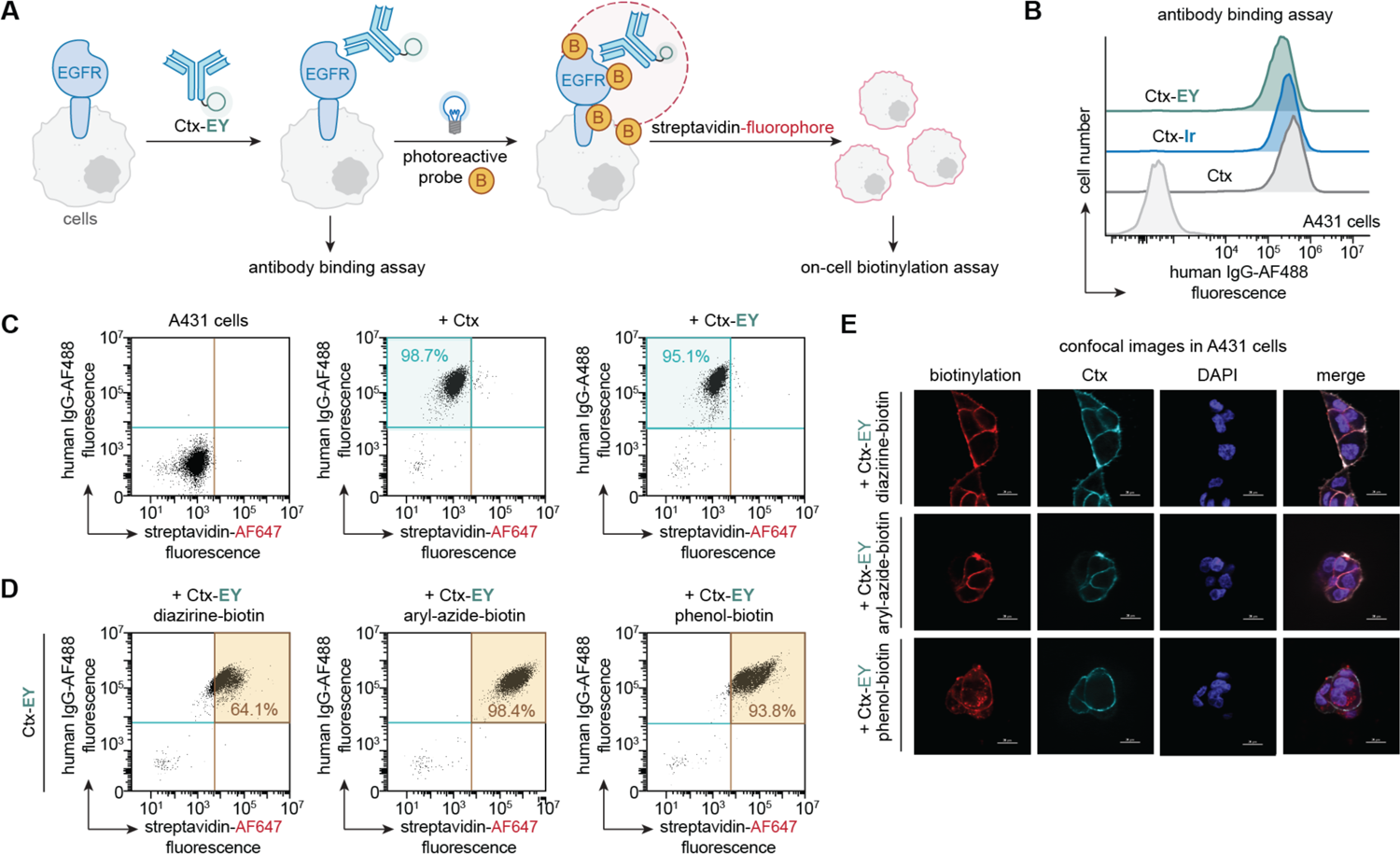
Ctx-EY enables EGFR-dependent labeling on cells with different photo-probes. **(A)** General on-cell labeling workflow using Ctx-EY conjugate and detection of biotin labeling using fluorescent streptavidin-AF647. **(B)** Cellular binding assay of 100 nM Ctx, Ctx-EY or Ctx-Ir conjugates on A431 cells via flow cytometry analysis shows similar on-cell binding. **(C, D)** Quantitative on-cell binding **(C)** and on-cell labeling **(D)** with diazirine-, aryl-azide- and phenol-biotin triggered by 100 nM Ctx-EY on A431 cells via flow cytometry analysis. **(E)** Confocal microscopy imaging of antibody binding and on-cell biotinylation of Ctx-EY on A431 cells shows labeling mostly confined to cell surface. Scale bar=20 µm.

We next performed on-cell proximity labeling with diazirine-, aryl-azide- and phenol-biotin photo-probes upon blue LED illumination (**Fig. 3D** and **fig S5A-D**). We tested a range of Ctx-EY concentrations and observed efficient biotinylation on cells at 100 nM (**Fig. 3D** and **fig. S5A**). The diazirine-biotin, aryl-azide biotin, and phenol-biotin labeling caused a major shift of biotinylation in the flow cytometry profile of 64%, 98%, and 94%, respectively. This is consistent with the order of labeling efficiencies observed *in vitro*. The Ctx-Ir only activated cell biotinylation with diazirine-biotin and aryl-azide-biotin and not phenol-biotin (**fig. S5B**).

We further visualized cell biotinylation induced by Ctx-EY via confocal microscopy (**Fig. 3E** and **fig. S5E-F**). The labeled A431, A549 and NCI-H441 cells were co-stained with both anti-human IgG-AlexFluor488 and streptavidin-AlexaFluor647 to visualize Ctx and biotinylation, respectively. We confirmed that the Ctx-EY conjugate was located on the cell membrane. Biotinylation using the diazirine- and aryl-azide-biotin photo-probes were observed mainly on the cell membrane, whereas the phenol-biotin labeling was more diffuse, consistent with the longer half-life and labeling range of the phenoxyl radical.

We next developed a proteomics workflow to label the EGFR neighborhood (**Fig 4A**), focusing first on A431 cells with highest levels of EGFR and using the most reactive diazirine-biotin photo-probe. We incubated A431 cells with or without EGF competition first and then performed the on-cell biotinylation workflow using Ctx-EY, followed by biotin enrichment using neutravidin beads. WB analysis confirmed selective biotinylation of EGFR which was ablated in the presence of EGF (**Fig. 4B**). We also observed dose-dependent EGFR labeling over a wide range of Ctx-EY concentrations of 1-1000 nM, which was competed off by either EGF or unlabeled Ctx (**fig. S6A**).

**Fig. 4.**
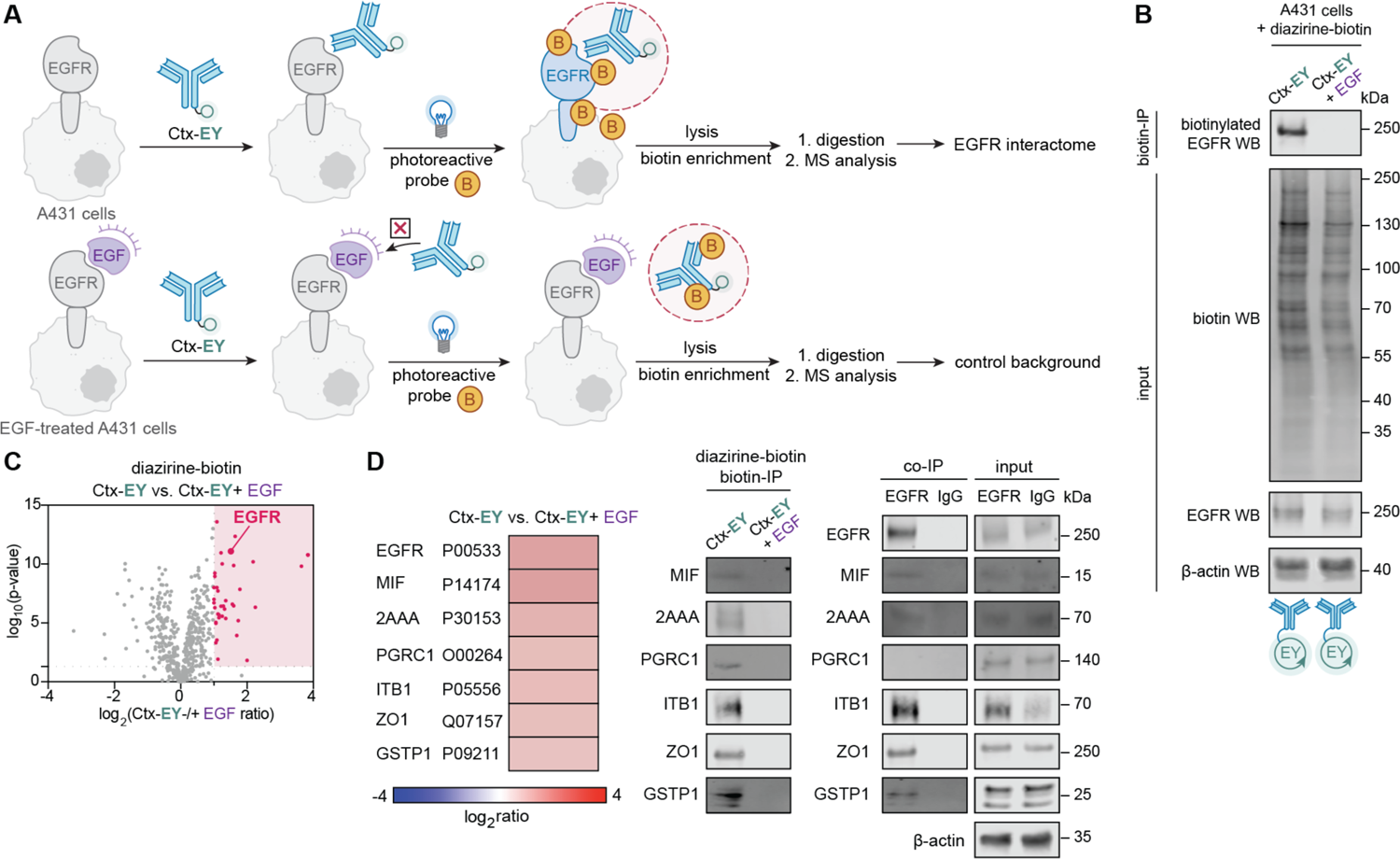
High-resolution profiling of the EGFR neighborhood using MultiMap. **(A)** General proteomics workflow of interactome profiling using Ctx-EY conjugate with or without EGF competition. **(B)** WB showing biotinylation using the diazirine-biotin photo-probe on A431 cells using Ctx-EY in the absence and presence of EGF competition. **(C)** Volcano plot of Ctx-EY-mediated labeling of EGFR with or without EGF on A431 cells using diazirine-biotin. 41 significantly enriched proteins (log_2_(ratio) ≥ 1, p-value<0.05, unique peptide ≥ 2, n=3) are highlighted in red and listed in **Table S9**. **(D)** Enrichment ratios for six top protein hits are displayed. Full heatmap is presented in **fig. S6B** and **Table S9**. All six proteins were confirmed to be biotinylated using biotin-IP blots, and five were selectively enriched in a separate EGFR co-IP experiment. All immunoblot images are representative of at least two biological replicates.

Cells were treated with Ctx-EY in the presence or absence of EGF competition and biotinylated proteins were captured on neutravidin beads and digested on-bead with trypsin. Samples were prepared in biological triplicate for MS analysis using label-free quantitation (volcano plot shown in **Fig. 4C**, heatmap shown in **Fig. S6B**). We identified a total of 536 proteins with 41 proteins enriched by more than two-fold with Ctx-EY relative to EGF competition (log_2_(ratio)≥1, p-value<0.05, unique peptide≥2; **Table S9** and **Fig. S6C-D**). EGFR was among the highly enriched as expected. Gene Ontology (GO) analysis showed a significant representation of biological processes that include regulation of phosphatase activity as well as molecular function entities such as phosphatase activator activity (**fig. S6D**). These features are consistent with the functional roles of EGFR signaling and suggest that the enriched EGFR interactors are accurately represented.

We orthogonally confirmed that six top hits were biotinylated by biotin-IP, where streptavidin pull-down samples were analyzed by WB using specific antibodies following proximity labeling (**Fig. 4D**). Among them, five were observed to co-IP with EGFR (**Fig. 4D**). All six proteins are known to either functionally interact with EGFR or found in immunoprecipitation experiments (*38–40*). These include ITB1, which is critical for stable maintenance for EGFR on the cell membrane (*41, 42*), as well as macrophage migration inhibitory factor (MIF), an immunostimulatory cytokine regulated by matrix metalloproteinase 13 (MMP13) known to be inhibitory for EGFR activation (*43*). Others include substrates of EGFR such as glutathione S-transferase P1 GSTP1 (*44*) and tight junction protein ZO1 (*45*), both of which are known to be activated upon phosphorylation by EGFR. One target membrane-associated progesterone receptor component 1, PGRC1, was not observed in EGFR co-IP experiment, and we would expect that some interactions may not be strong enough to survive the co-IP workup in these cells.

### Multi-scale EGFR interactome profiled via MultiMap

Having demonstrated the proteomic workflow of Ctx-EY triggered biotinylation on cells expressing high levels of EGFR, we expanded to cells expressing modest levels of EGFR. Lung cancer cell line, A549, for example, express lower amounts of EGFR (nTPM=59.7), which is more typical of native membrane proteins (*37*). We applied all photo-probes and showed EGFR was selectively biotinylated with each probe (**fig. S7A**). Applying the proteomics workflow, we then identified EGFR neighbors enriched with diazirine-biotin, aryl-azide-biotin and phenol-biotin by comparing labeling with Ctx-EY in the absence and presence of EGF (**Fig. 5A**). We found that EGFR is one of the most enriched proteins from all three datasets (**Table S10-12**). Enriched proteins were identified with the same statistical thresholds [log_2_(ratio)≥1, p-value<0.05, unique peptide≥2], allowing us to compare protein identities across reactions with different photo-probes. We identified 72 proteins using diazirine-biotin, 188 using aryl-azide-biotin, and 188 using phenol-biotin (**Tables S10-12** and **fig. S7B-D**).

**Fig. 5.**
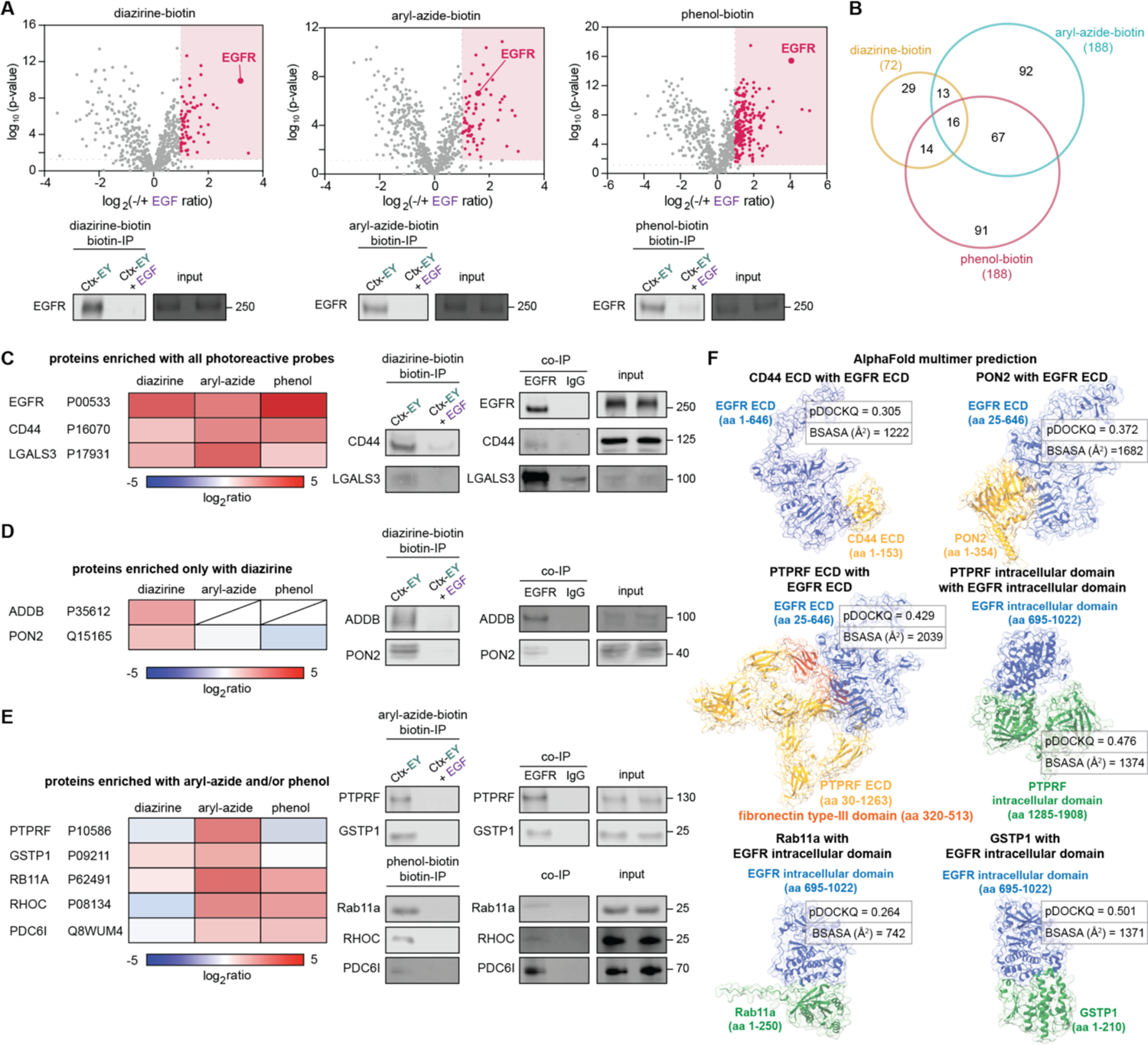
MultiMap reveals a multi-scale EGFR interactome network. **(A)** Volcano plots of Ctx-EY mediated EGFR interactome profiling on A549 cells using three different photo-probes (biotin-diazirine, aryl-azide-biotin, or phenol-biotin, respectively, n=3). Significantly enriched proteins (log_2_(ratio)≥1, p-value<0.05, unique peptide≥2) are highlighted in red and listed in **Table S10-12**. **(B)** Venn diagram of EGFR interactome enriched from A431 cells using different photo-probes. **(C)** Enrichment ratios and validation of protein hits using all three photo-probes. **(D)** Enrichment ratios and validation of protein hits from only the diazirine-biotin dataset. **(E)** Enrichment ratios and validation of protein hits from both aryl-azide-biotin and/or phenol-biotin datasets. **(F)** AlphaFold-Multimer predictions of EGFR complexes confirmed the direct interactions of EGFR with interactors found via MultiMap. EGFR ECD or ICD (blue) is shown in complex with the corresponding interactor protein (yellow or orange-yellow for the ones interacting with EGFR ECD, green for the ones interacting with EGFR ICD) along with the pDockQ scores and BSASA. All immunoblot images are representative of at least two biological replicates.

As represented in a Venn diagram (**Fig. 5B**), there were a total of 322 unique proteins enriched over the controls in at least one of the three photo-probes. The aryl-azide-biotin and phenol-biotin labeled more proteins than diazirine-biotin reflecting their higher yields and their relatively long labeling radii. We found that >80% of the enriched proteins were annotated as plasma membrane proteins in UniProt (plasma membrane, GO:0005886) for all three photo-probes. GO enrichment analysis suggested molecular functions such as EGFR activity and EGF binding were highly enriched (**fig. S7D**).

Sixteen candidate neighbors were identified in all three datasets of MultiMap (**Fig. 5B**). While no direct structural evidence has been reported for EGFR with any of these, CD44 and Galectin-3 have been functionally associated with EGFR: CD44 regulates EGFR functions in the presence of CD147 and hyaluronan (*46, 47*); Galectin-3 regulates EGFR localization and its interactions suggested through genetic studies in pancreatic cancers (*48*). Both targets were further validated by biotin-IP and EGFR co-IP (**Fig. 5C**), supporting that they are proximal neighbors of EGFR.

We next expanded our list to the 29 proteins that were in common for diazirine-biotin and aryl-azide-biotin, the photo-probes with high labeling resolutions (**Fig. 5B**). Among them, we found a paraoxonase, PON2, as well as two proteins associated with RTK phosphorylation and activation: beta-adducin ADDB and MAP kinase pathway member BRAF (*49, 50*). Both PON2 and ADDB were detected by biotin-IP and EGFR co-IP. Interestingly, BRAF, a cytosolic protein was enriched by biotin-IP but not EGFR co-IP suggesting it is close but may not be in physical contact (**Fig. 5D** and **fig. S7E**). Between the diazirine and aryl-azide datasets, we identified the known EGFR functional interactors such as Tid1 (*51*) and ITB1 also found in the A431 cell experiments (33, 34), as well as previously unreported interaction partners including CKAP4 and RAC1 (**fig. S7E**).

The aryl-azide-biotin and phenol-biotin experiments contributed more proteins (293 in total). Among the top hits were the tyrosine-protein phosphatase receptor, PTPRF, glutathione transferase GSTP1, small GTPase Rab11a, Rho-related GTP binding protein RHOC and ESCRT protein PDC6I (**Fig. 5E** and **fig. S7E**). Remarkably, all were detected by biotin-IP with ten out of eleven of these proteins co-IPed with EGFR, suggesting that they form relatively stable complexes. To provide a structural level of analysis, we turned to AlphaFold-Multimer, an exciting extension of AlphaFold developed over the last few years that uses artificial intelligence to generate plausible models of binary protein complexes (*24, 52, 53*). This community has developed scoring metrics such a predicted DockQ score (pDockQ), where a threshold of >0.23 retrieves 51% of true-positive interacting proteins with a false-positive rate of ∼1% in large test set models (*54*). Additional criteria can be applied including buried solvent accessible surface area (BSASA)≥500Å^2^ (*55, 56*), predicted local distance difference test (pLDDT)>50 for the interface residues and minimum predicted alignment error (PAE)<15 Å as described previously (*52*). As a true positive example, we derived an AlphaFold-Multimer model of the EGF:EGFR complex (**fig. S8A** and **Table S13)** that closely overlaid that of the known structure of EGF:EGFR (PDB: 1IVO, RMSD between 469 atom pairs is 0.924 Å) (*57*).

We applied AlphaFold-Multimer to candidate neighbors validated by biotin-IP and/or EGFR co-IP in A431 and A549 cells and generated a total of 29 models. As shown in waterfall plots, the average pDockQ score (0.298) and BSASA (1466Å^2^) for the 29 EGFR-protein pairs were both above the established criteria suggesting direct interactions (**fig. S8B** and **Table S13**). We next applied AlphaFold-Multimer to compute models of all potential heterodimeric complexes from **Fig. 4C** and **Fig. 5A** (**Table S13**). To increase the accuracy of the models for transmembrane proteins (*58*), we calculated separately the ECD (aa 1-646) and intracellular domain (ICD, aa 695-1022) of EGFR and paired them with the corresponding ECDs or ICDs of transmembrane protein targets. As previously described, we retained only high-confidence AlphaFold-Multimer models [average pLDDT>50, minimum predicted Alignment Error (PAE)<15Å] (*52*) and performed further filtering using the aforementioned criteria [pDockQ score≥0.23, BSASA≥500 Å^2^]. The final list of validated complexes included the binary complexes of EGFR ECD with CD44 ECD (aa 1-153, pDockQ=0.375), PON2 (pDockQ=0.372) and MIF (aa 1-115, pDockQ=0.375) (**Fig. 5F** and **fig. S8C-F**). In addition, the ICD of EGFR is predicted to bind Rab11a (pDockQ=0.264), GSTP1 (pDockQ=0.535) and RAC1 (pDockQ=0.387) (**Fig. 5F** and **fig. S9A-C**). Most interestingly, one of the AlphaFold-Multimer complexes predicted with the highest confidence is a cell-surface phosphatase PTPRF, where PTPRF ECD binds EGFR ECD (pDockQ=0.429) and likewise, the PTPRF ICD binds the EGFR ICD (pDockQ=0.476) (**Fig. 5F** down).

### MultiMap can capture distal synaptic protein networks

Extracellular protein-protein interactions occur not only *in cis* on the cell membrane but also *in trans* between cell-cell junctions. To explore PLP of cell synapses using MultiMap at different labeling radii, we assembled a co-culture system where the cell-cell interaction was induced by a bispecific T cell engager (BiTE) (**Fig. 6A**). This BiTE contained the Ctx Fab genetically fused to an α-CD3 scFv (OKT3) (*59*). Two different cells were utilized in the co-culture system: a HEK293T cell was engineered in house to overexpress a Flag-tagged-EGFR (HEK-Flag-EGFR) and well-established Jurkat cells expressing a NFAT-GFP reporter (*60*). In this design, the Flag tag served as an orthogonal ecto-epitope for an EY-conjugated α-Flag nanobody (α-Flag-EY), thus allowing selectively recognition of Flag-tagged EGFR. In order to separately characterize the labeling on HEK-Flag-EGFR and Jurkat NFAT-GFP cells, we used α-CD3-PE signal to allow facile separation of CD3^+^ Jurkat cells from CD3^-^ HEK-Flag-EGFR. Levels of cis- and trans-labeling from α-Flag-EY were determined by flow cytometry. Proteins labeled with different photo-probes were enriched using streptavidin beads and analyzed by WB (**Fig. 6A**).

**Fig. 6.**
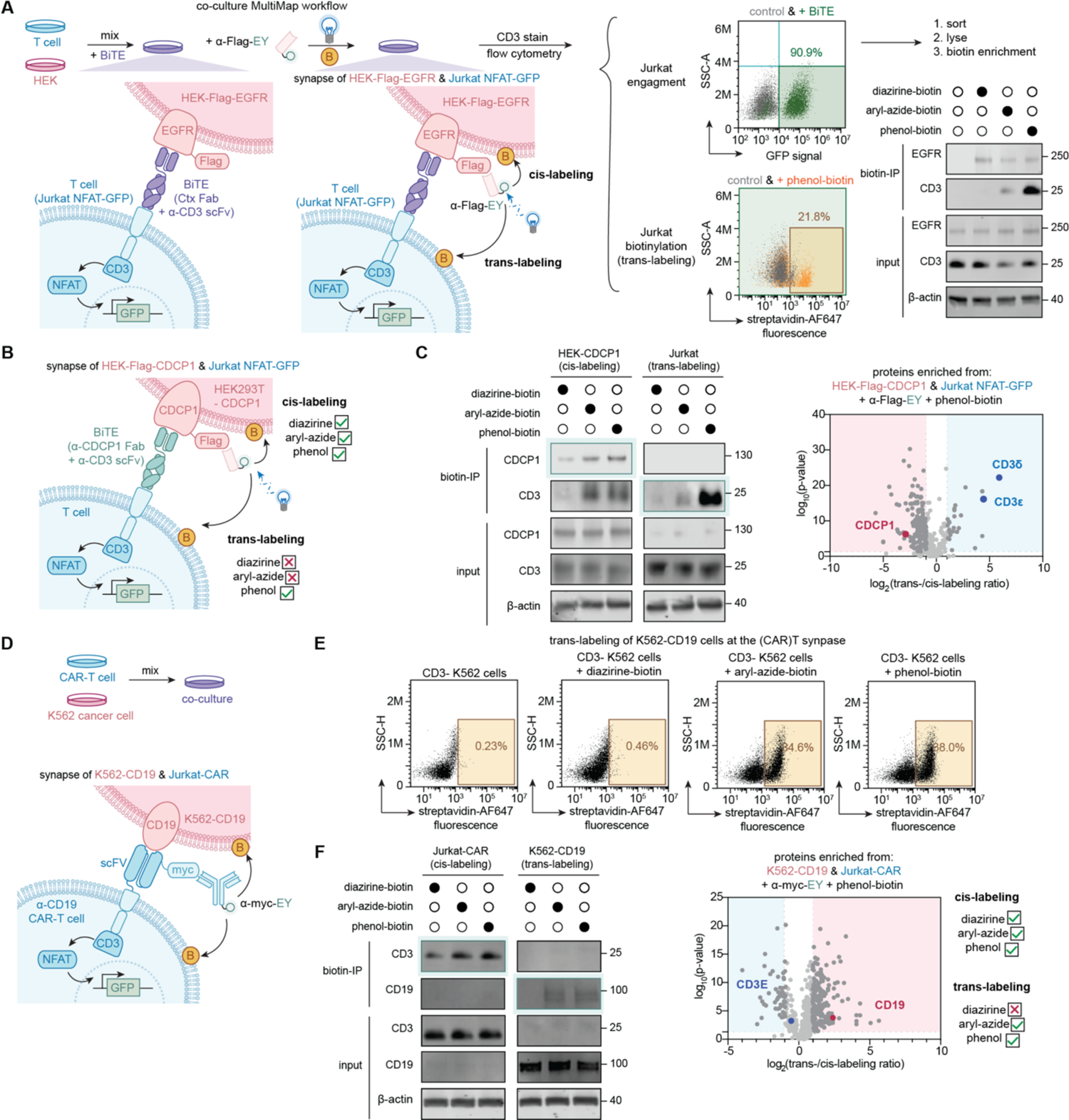
MultiMap enables targeted snapshots of local cell-cell synapses. **(A)** On-cell labeling of the T-cell synapse using a bispecific T cell engager (BiTE) that recognizes EGFR. Jurkat NFAT-GFP and HEK293T-Flag-EGFR were co-cultured in the presence of the BiTE before MultiMap was performed using an EY-conjugated α-Flag nanobody (α-Flag-EY). Cell-cell engagement was monitored by NFAT-GFP reporter gene activation. Photocatalytic labeling was characterized by flow cytometry before biotin-enriched proteins were analyzed by WB. **(B)** Target biotinylation of CDCP1 and CD3 at the T cell synapse using a bispecific T cell engager (BiTE) that recognizes CDCP1. Longer labeling radius using phenol-biotin was necessary for trans-labeling on Jurkat NFAT-GFP. **(C)** Volcano plot of proteins biotinylated on HEK-Flag-CDCP1 (cis-labeling) and Jurkat NFAT-GFP (trans-labeling) using phenol-biotin (n=3). Significantly enriched proteins from HEK-Flag-CDCP1 (log_2_(ratio) ≤ **-**1, p-value<0.05, unique peptide ≥ 2) or Jurkat NFAT-GFP (log_2_(ratio)≥1, p-value<0.05, unique peptide≥2) are highlighted in blue and red, respectively. Full protein lists were shown in **Table S14**. **(D)** Scheme of on-cell labeling of anti-CD19 chimeric antigen receptor (CAR)T cell system. **(E) (**CAR)T cell-mediated trans-labeling of K562 cancer cells using all photo-probes. **(F)** Target biotinylation of CD3 and CD19 at the CAR-T synapses using WB and MS analysis. Cells were sorted to differentiate cis- and trans-labeling before biotinylated proteins were enriched for analysis. Both phenol-biotin and aryl-azide-biotin enabled trans-labeling. Volcano plot of proteins biotinylated on Jurkat-CAR (cis-labeling) and K562-CD19 (trans-labeling) using phenol-biotin (n=3). Significantly enriched proteins from Jurkat-CAR (log_2_(ratio)≤**-**1, p-value<0.05, unique peptide≥2) or K562-CD19 (log_2_(ratio)≥1, p-value<0.05, unique peptide≥2) are highlighted in blue and red, respectively. Full protein lists were shown in **Table S15**. All immunoblot images are representative of at least two biological replicates.

We first monitored BiTE engagement between HEK-Flag-EGFR and Jurkat NFAT-GFP cells using the standard GFP reporter gene readout. As expected, we observed dose-dependent BiTE activation of cell-cell engagement, with the GFP signal shifted to 80.3% in the presence of 8 nM EGFR BiTE and 92.3% with 50 nM BiTE (**Fig. S10A**). The GFP signal shift was not affected by the presence of α-Flag-EY, indicating that the Flag tag recognition did not interfere with the cell synapse engagement. We then performed the MultiMap workflow using four photo-probes of increasing labeling range: diazirine-biotin, aryl-azide-biotin, biocytin-hydrazide and phenol-biotin. We monitored biotinylation *in cis* for HEK-Flag-EGFR and *in trans* for Jurkat NFAT-GFP using a streptavidin-AlexaFluor647 signal (**Fig. 6A** and **fig. S10B-C**). Cis-labeling of HEK-Flag-EGFR cells occurred for >60% of cells for the diazirin-biotin, aryl-azide-biotin and phenol-biotin with ∼29% for the biocytin hydrazide (**fig. S10B**). In sharp contrast, minimal shift (∼3-4%) was observed on control HEK-EGFR cells without the Flag tag, suggesting that α-Flag-EY is selectively recognizing the Flag tag. Interestingly, the trans-labeling of the Jurkat cells using the shorter-range diazirine-biotin, aryl-azide-biotin was limited to 3-4% (**fig. S10C**), while the intermediate-range biocytin-hydrazide and long-range phenol-biotin labeled 9% and 22%, respectively) (**Fig. 6A** and **fig. S10C**). This is consistent with the cell-cell synapse distance based on the length of the Fab-ScFv BiTE (*61*), plus the size of the EGFR ECD and the CD3 complex. By further analysis via WB, the cis-target EGFR was observed enriched by biotin-IP in the presence of the three photo-probes, whereas trans-target CD3 was only significantly enriched in the phenol-biotin sample, with moderate amount observed in the aryl-azide-biotin-labeled sample (**Fig. 6A**). Thus, longer-range photo-probes are more efficient for trans-labeling.

To further expand the generality of MultiMap for cell-cell synapses, we tested the BiTE system to two other cancer targets, HER2 and CDCP1 (**Fig. 6B** and **fig. S11A-E**). We fused α-HER2 Fab sequence (Trz Fab, **fig. S10A-C**) and a previously generated α-CDCP1 Fab (4A06) (**Fig. 6B-C** and **fig. S11D-E**) (*59*) onto the CD3 scFv scaffold. Again, we observed dose-dependent cell-cell engagement in the presence of the engineered BiTEs and antigen-expressing cells (**fig. S11B** and **fig. S11D**). On-cell biotinylation results were similar to the EGFR BiTE system; cis-labeling was found with all three photo-probes and trans-labeling activated primarily with phenol-biotin (**fig. S10C** and **fig. S11E**). In particular, by separating HEK293T-CDCP1 and Jurkat-NFAT-GFP cells, we confirmed selective biotinylation of CDCP1 with all three photo-probes, and CD3 only with phenol-biotin (**Fig. 6B**). We quantitatively profiled the proteins captured at the cell synapse of HEK-Flag-CDCP1 and Jurkat NFAT-GFP in biological triplicate (**Fig. 6C**, **Table S14**). We discovered that CDCP1 was enriched in the cis-labeled samples. Proteins from the CD3 complex including CD3δ and CD3ε were highly enriched in the trans-labeled samples. This observation indicates that interactome in the cell-cell synapse can be captured by the MultiMap workflow.

Lastly, we evaluated MultiMap labeling at a (CAR)T cell-cell synapse (**Fig. 6D** and **fig. S11F-G**). Jurkat cells expressing a Myc-tagged CAR construct (Jurkat-CAR) that targets CD19 were mixed with K562 cancer cells expressing CD19 ectodomain (K562-CD19). With an EY-conjugated α-myc antibody (α-myc-EY), we performed our workflow by introducing cis-labeling on K562-CD19 and trans-labeling on Jurkat-CAR cells. We confirmed cell engagement by monitoring the CAR activation with or without K562 cells (**fig. S11F**). We observed cis-labeling on interacting CAR cells with all photo-probes (**fig. S11G**). On the other hand, both aryl-azide-biotin and phenol-biotin achieved trans-labeling, with much lower level of biotinylation using the short-range diazirine-biotin (**Fig. 6E**). The same results were confirmed by WB analysis (**Fig. 6F**). These results are in line with the estimate of cell-cell distance between CAR-induced synapse at ∼120 Å according to AlphaFold prediction (*62*), which is shorter than BiTE-induced synapse. We finally sorted each cell type for proteomics analysis and found both CD19 and CD3 component enriched for trans-labeling and cis-labeling (**Fig. 6F** and **Tables S15**). Taken together, our data suggests that MultiMap can label cells at the cell-cell synapses and map the proteins in proximity via PLP. Only minimal alternation to the existing workflow is needed for labeling in different cell-cell engagement scenarios. Looking forward, we anticipate that this platform will be a useful technology to identify key proteins in different synaptic environments.

## Discussion

Here we demonstrate a multi-scale PLP technology, MultiMap, that enables proximity labeling and interactome profiling with adjustable resolution depending on the half-life of the photo-probe using a single photocatalyst EY. EY is a unique photocatalyst in its ability to trigger a broad range of photo-probes. It is commercially available, bio-compatible, and shown to be readily conjugated to seven different proteins and antibodies by commonly accessible methods. Simple targeting by EY-conjugated antibodies obviates the need for cell engineering. EY-mediated labeling is rapid and light-dependent, which potentially allows kinetic control of the labeling. Future structure-activity relationship studies on EY, as done for rhodamine- or fluorescein-based scaffolds (*63*), could enhance our understanding of the photochemical mechanisms of EY, as well as improve its spectral properties, activation efficiency, and use in tissues (*20, 27*).

Coupled with standard biochemical validation and the recently developed AlphaFold-Multimer algorithm for structural prediction (*24*), the MultiMap proximity labeling proteomics workflow provides three orthogonal and integrated pillars for high-resolution profiling of protein neighborhoods. In addition to identification of new potential neighbors, we detected many proteins known to functionally interact with EGFR that are reported to stabilize, modulate, or act as substrates for EGFR. One of the most striking targets identified was the phosphatase, PTPRF, which could be a functional off-switch for EGFR. Interestingly, AlphaFold-Multimer predicts the ECD of PTPRF binds the back side of the EGFR ECD away from the dimer interface, and that the ICD of the phosphatase binds to the intracellular kinase domain of EGFR. Despite the fact that EY-antibodies recognize extracellular targets, we found that some of the high-confidence hits were intracellular proteins. Some are known to functionally associate with EGFR, for which high-confidence AlphaFold-Multimer binary models were constructed. It is not impossible that the labeling is caused by cell penetrance of the activated photo-probe when triggered by EY. We believe future work could fine-tune the properties of photo-probes such as charge and hydrophobicity to achieve extracellular-only or organelle-specific labeling.

It is unlikely that all these identified neighbors bind simultaneously to EGFR. In fact, some proteins are predicted to bind over the same sites. These data suggest EGFR can be in multiple neighborhoods which are dynamic and may have multiple functions yet to be revealed. It is also important to note that the binder we used in this study, Ctx, is an inhibitor of EGFR function. Thus, candidates identified in our studies are specifically from the EGFR off-state neighborhood. We envision that MultiMap will be useful to study on-state, drug-bound, and resistance mutant neighborhoods, which will give a comprehensive map of the EGFR interactomes.

MultiMap was also effective for long-range labeling of cell-cell synapses. As shown for the ones activated by BiTE or (CAR)T, we found that spatial variability among synaptic junctions can be addressed by using photo-probes with different labeling radii. The unique advantage of MultiMap allowing multi-scale labeling, potentiates its application for interactome profiling of additional intercellular interaction networks. In cases where antibodies are not available, one can use a genetically encoded tag on the target ECD, similar to the Flag and myc ecto-tags we introduced in our study. We can also envision proteome-wide interactome profiling for membrane proteins using these ecto-tags on par with the scale for the intracellular OpenCell system (*64*).

Lastly, we recognize that despite the confidence and accelerated process in target identification via MultiMap, information on candidate neighborhoods warrants further confirmation via more in-depth structural, mutational, and functional studies. Nonetheless, we believe that MultiMap proximity labeling proteomics integrated with *in silico* prediction can begin to provide plausible models for binary protein interactions with high structural precision. This would be crucial step forward to begin to construct a structural map of the protein-protein interactome on the cell surface. In addition, understanding how the surfaceome functions in concert with the external environment communication may suggest new neo-complexes to target for both small molecules and biologics.

## Acknowledgements

We thank Jie Zhou, Kevin Leung, Thomas Bartholow, Johnathan Maza, Kaan Kumru, Corleone Delaveris, Paul Burroughs and James Byrnes for insightful discussions. We also thank Susanna Elledge for the Her2 Fab expression plasmids, Jhoely Duque-Jimenez and Xin Zhou from Harvard Medical School for CAR-T and K562-CD19 cells as well as Juntao Yu from Harvard Medical School for guidance in conducting the AlphaFold-Multimer analysis.

## Funding

We are grateful to generous support from NIH (1R01CA248323-01). K.S. is a Merck fellow of the Helen Hay Whitney Foundation. Z.Y. is supported by an NIH National Institute of General Medical Sciences F32 grant (1F32GM149084-01). A.S. is supported by NIH (P41GM109824 and R01GM083960). A.P. is supported by NIH (U19AI135990). The HDFCC LCA for cell sorting is funded by NIH (P30CA082103).

## Author contributions

Z.L. and J.A.W designed the project, analyzed the data and wrote the manuscript with input from all authors. K.S. and Z.Y. generated plasmids and purified proteins. I.L. assisted the cell sorting experiments. A.F. and D.L.S. assisted the mass spectrometry experiments. A.P. and A.S. assisted AlphaFold-Multimer analysis and BSASA calculation.

## Declaration of interests

The authors declare the following competing financial interest: J.A.W and Z.L filed a provisional patent on the multi-scale interactome profiling of membrane proteins using photocatalytic proximity labeling.

## Data and materials availability

All data are available in the main text or the supplementary materials.

## Notes

### Competing Interest Statement

The authors have declared no competing interest.

