## Supplemental information for "Multi-scale photocatalytic proximity labeling reveals cell surface neighbors on and between cells"

### Index

**Supplementary Figures:**
Fig. S1. Eosin Y (EY) as an organic photocatalyst that triggers the activation of photo-probes.
Fig. S2. Conjugation chemistry of EY onto BSA and Ctx for targeted protein biotinylation.
Fig. S3. Identification of specific sites and residues labeled with different biotin-photoprobes using EY-conjugated antibodies.
Fig. S4. Flow cytometry experiments with cells containing different levels of EGFR shows Ctx-EY binds to cells similarly to unconjugated Ctx.
Fig. S5. Ctx-EY can activate cell-surface biotinylation on A431, A549 and NCI-H441 cells using different photo-probes.
Fig. S6. Specific labeling of EGFR labeling on A431 cells by Ctx-EY.
Fig. S7. EGFR interactome profiling using Ctx-EY on live A549 cells.
Fig. S8. AlphaFold-Multimer predictions of EGFR binary complexes with EGFR interactors from MultiMap datasets.
Fig. S9. Additional predicted binary complexes of EGFR interactors with EGFR.
Fig. S10. Targeted labeling of BiTE-induced synapses between Jurkat NFAT-GFP and HEK-Flag-EGFR using α-Flag-EY.
Fig. S11. Extended applications of targeted cell synapse labeling.

**Methods:**
General methods and instrumentation.
General chemical methods and instrumentation.
Reaction monitoring by LC-MS.
Antibodies and biological reagents.
Plasmid Construction.
Cell culture.
Mammalian protein expression.
General protocol for antibody conjugation with EY.
Western blot protocol.
General flow cytometry.
On-cell antibody binding and biotinylation assay.
Recombinant protein biotinylation assay.
General protocol for antibody-EY labeling on cells.
Sample preparation for LC-MS/MS analysis.
Proteomics analysis of digested peptide samples.
Analysis of proteomics dataset.
Immunoprecipitation assays (co-IP) in live cells.
AlphaFold-Multimer prediction and analysis.
Cell co-culture.
Software.
Statistical analysis.
**Synthetic procedures.**
Scheme 1. Synthesis of DBCO-PEG$_4$-EY (**6**).
Scheme 2. Synthesis of diazirine-PEG$_3$-biotin (**9**).
**References.**

### Supplementary Figures:

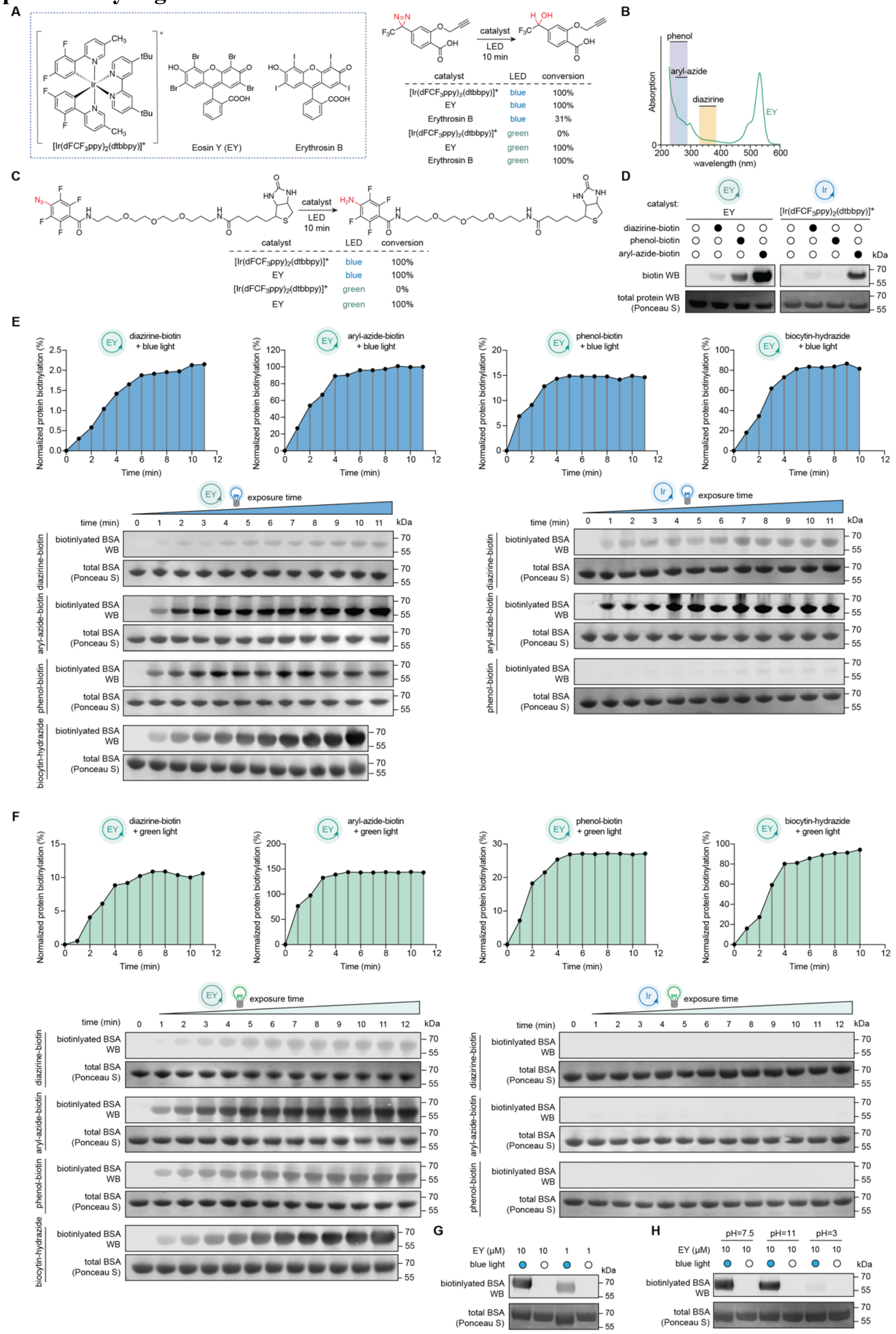

**Fig. S1. Eosin Y (EY) as an organic photocatalyst that triggers the activation of photo-probes.** **(A)** Eosin Y (EY) triggers conversion of diazirine in the presence of blue or green LED. Chemical structures of photocatalysts used in this study are shown. Conversion of the diazirine substrate was monitored via LC-MS. **(B)** Absorption peaks of the photoreactive warheads (diazirine, aryl-azide and phenol, shown in colored boxes) and EY. Photoreactive warheads cannot be activated by blue or green LED (*1, 2*) and thus require EY for activation. **(C)** EY triggers conversion of aryl-azide-biotin in the presence of blue or green LED. **(D)** EY triggers the biotinylation of bovine serum albumin (BSA) using diazirine-biotin, aryl-azide-biotin and phenol-biotin upon blue LED illumination. Quantification of biotinylation is shown in **Fig. 1C**. **(E)** Time-dependent biotinylation on BSA demonstrated the rapid kinetics for labeling using EY with blue LED activation. Biotinylation levels of BSA in were tracked via Western blot (WB) analysis and quantified, indicating that EY can trigger >90% conversion with 3 min illumination with blue LED. **(F)** Time-dependent biotinylation on BSA demonstrated the rapid kinetics for labeling using EY with green LED activation. Biotinylation levels of BSA were tracked via WB analysis and quantified. **(G)** EY-induced biotinylation using diazirine-biotin is dose-dependent. **(H)** Photocatalytic ability of EY was not affected in basic condition (pH=11), but was significantly inhibited in acidic condition (pH=3). All immunoblot images are representative of at least two biological replicates.

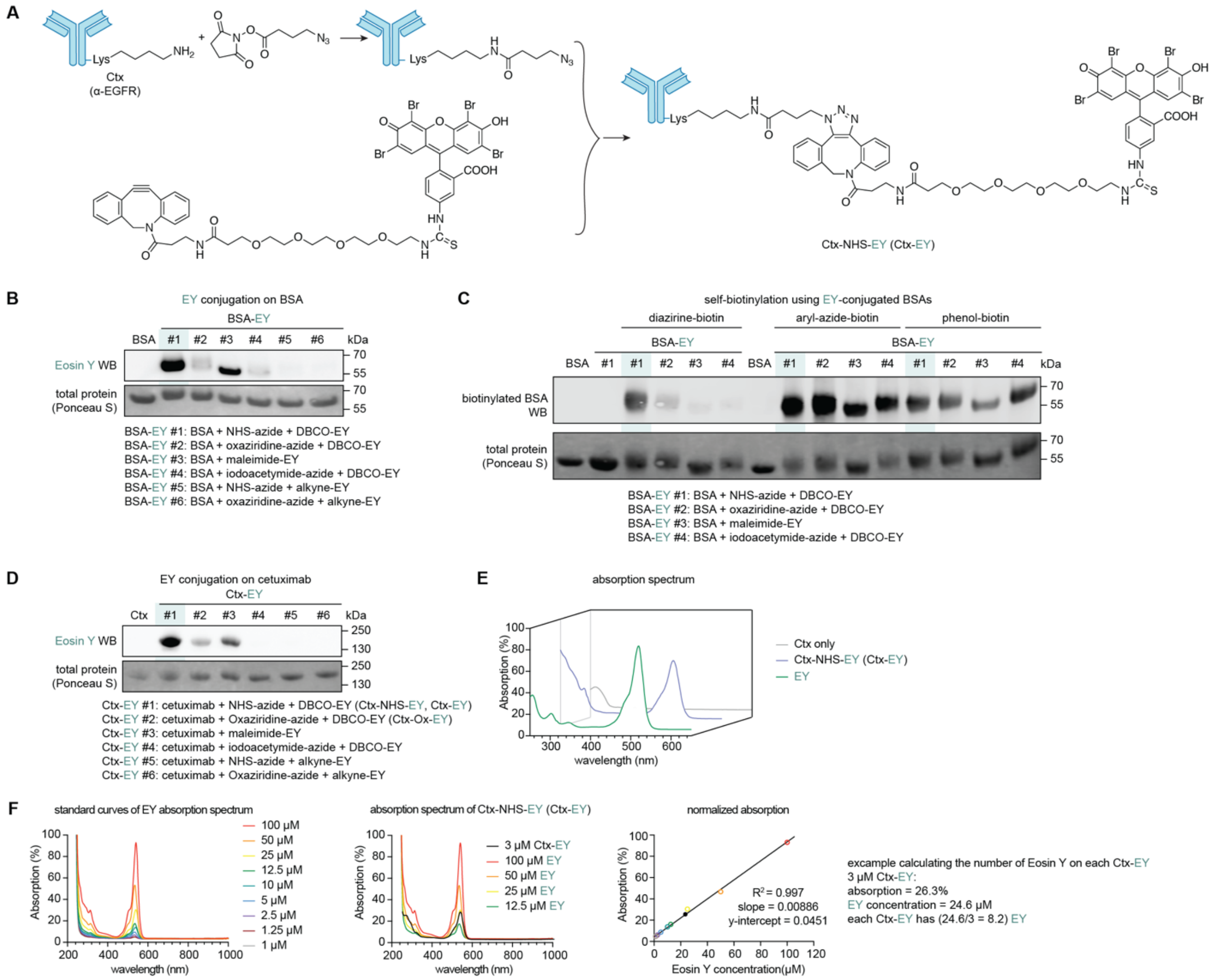


**Fig. S2. Conjugation chemistry of EY onto BSA and Ctx for targeted protein biotinylation.** **(A)** Chemical structure of Lys-specific conjugation of EY using DBCO-PEG$_4$-EY. Synthetic route of DBCO-PEG$_4$-EY is shown in **Scheme 1**. **(B)** Screening of conjugation methods on BSA using different covalent warheads: NHS, oxaziridine, maleimide and iodoacetamide. The NHS-based amine coupling (highlighted in green) produced the highest extent of conjugation owed to presence of available Lys residues. **(C)** Evaluation of self-biotinylation using different EY-conjugated BSA constructs. Three photo-probes (diazirine-, aryl-azide- and phenol-biotin) were tested. **(D)** Screening of conjugation methods on Ctx with different residue-specific labeling methods mirrors results seen with BSA in Panel (**B**). **(E)** Overlayed absorption spectra of Ctx, Ctx-EY and EY. Conjugation of EY onto Ctx has minimal effect on the EY's absorption spectrum. **(F)** Calculation of conjugated EY stoichiometry using the photochemical property of EY as a dye. An example of Ctx-EY is shown, demonstrating that an average of eight EY molecules was conjugated per Ctx. All immunoblot images are representative of at least two biological replicates.

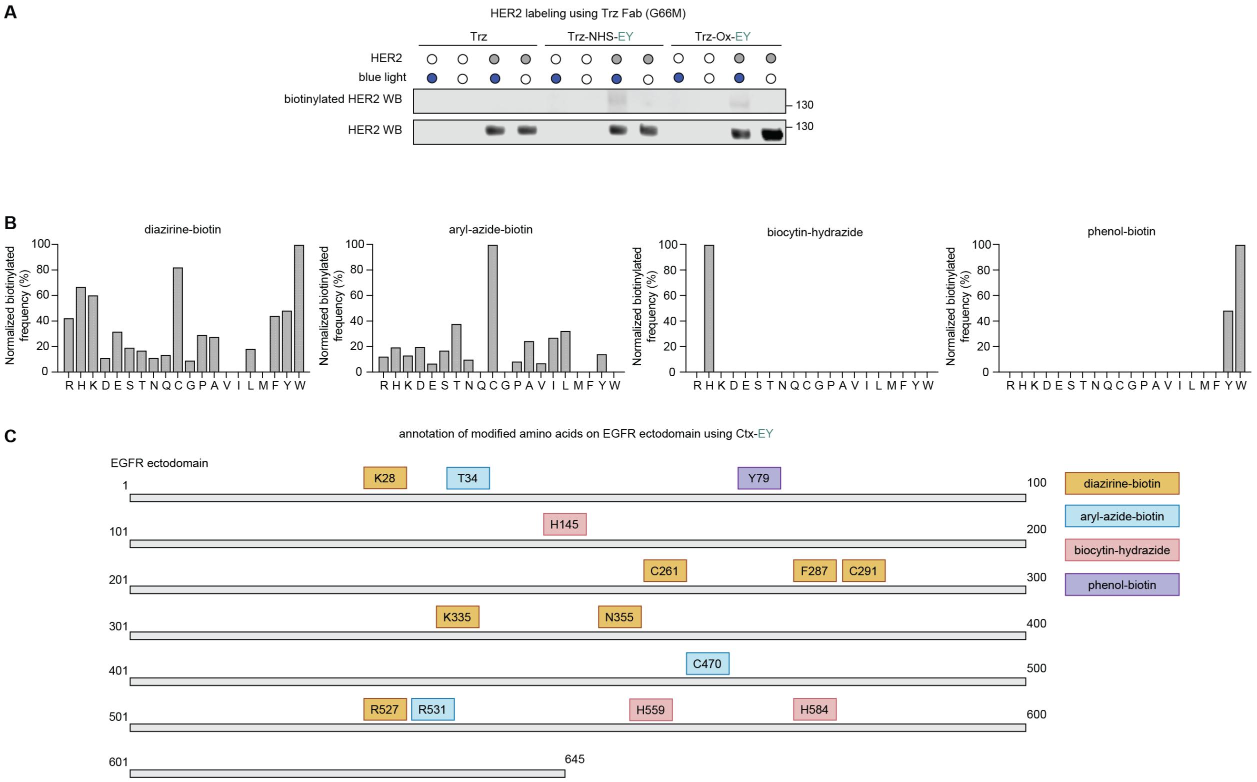


**Fig. S3. Identification of specific sites and residues labeled with different biotin-photoprobes using EY-conjugated antibodies. (A)** Selective labeling of the ecto-domain of HER2 using EY-conjugated Trz. A Trz Fab mutant (G68M) (*3*) was conjugated using covalent warheads of NHS or oxaziridine, enabling targeted labeling of the purified HER2 ectodomain (aa 23-652) *in vitro*. **(B)** Statistics of amino acid labeling preference with four photo-probes from combined datasets of observed biotinylated peptides on BSA (45 peptides in total), Ctx and EGFR ECD (88 peptides in total). Spectrum assignments were shown in **Table S1-8**. **(C)** Identification of the EGFR ectodomain residues that were modified with different photo-probes. Amino acids labeled with diazirine-biotin, aryl-azide-biotin, phenol-biotin or biocytin-hydrazide are mapped on the sequence of EGFR or the crystal structure of EGFR ectodomain (PDB: 1YY9) shown in **Fig. 2E**. Spectrum assignments were shown in **Table S5-8**. Immunoblot images are representative of at least two biological replicates.

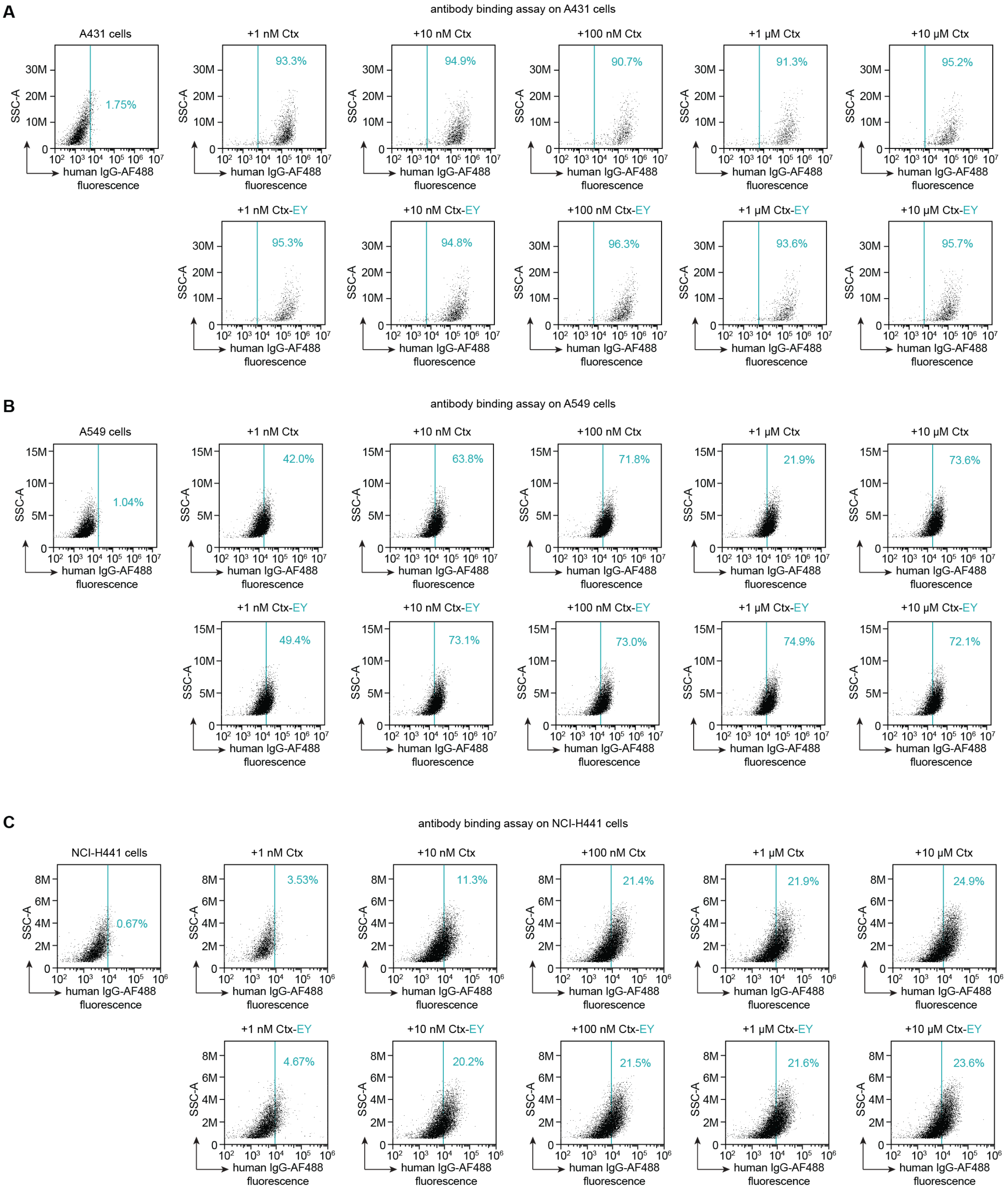


**Fig. S4. Flow cytometry experiments with cells containing different levels of EGFR shows Ctx-EY binds to cells similarly to unconjugated Ctx. (A)** Quantitative on-cell binding assay of Ctx and Ctx-EY on A431 cells with high EGFR expression levels (EGFR nTPM: 2978). **(B)** Quantitative on-cell binding assay of Ctx and Ctx-EY on A549 cells with low EGFR expression levels (EGFR nTPM: 59.7). **(C)** Quantitative on-cell binding assay of Ctx and Ctx-EY on NCI-H441 cells with very low EGFR expression levels (EGFR nTPM: 29.8).

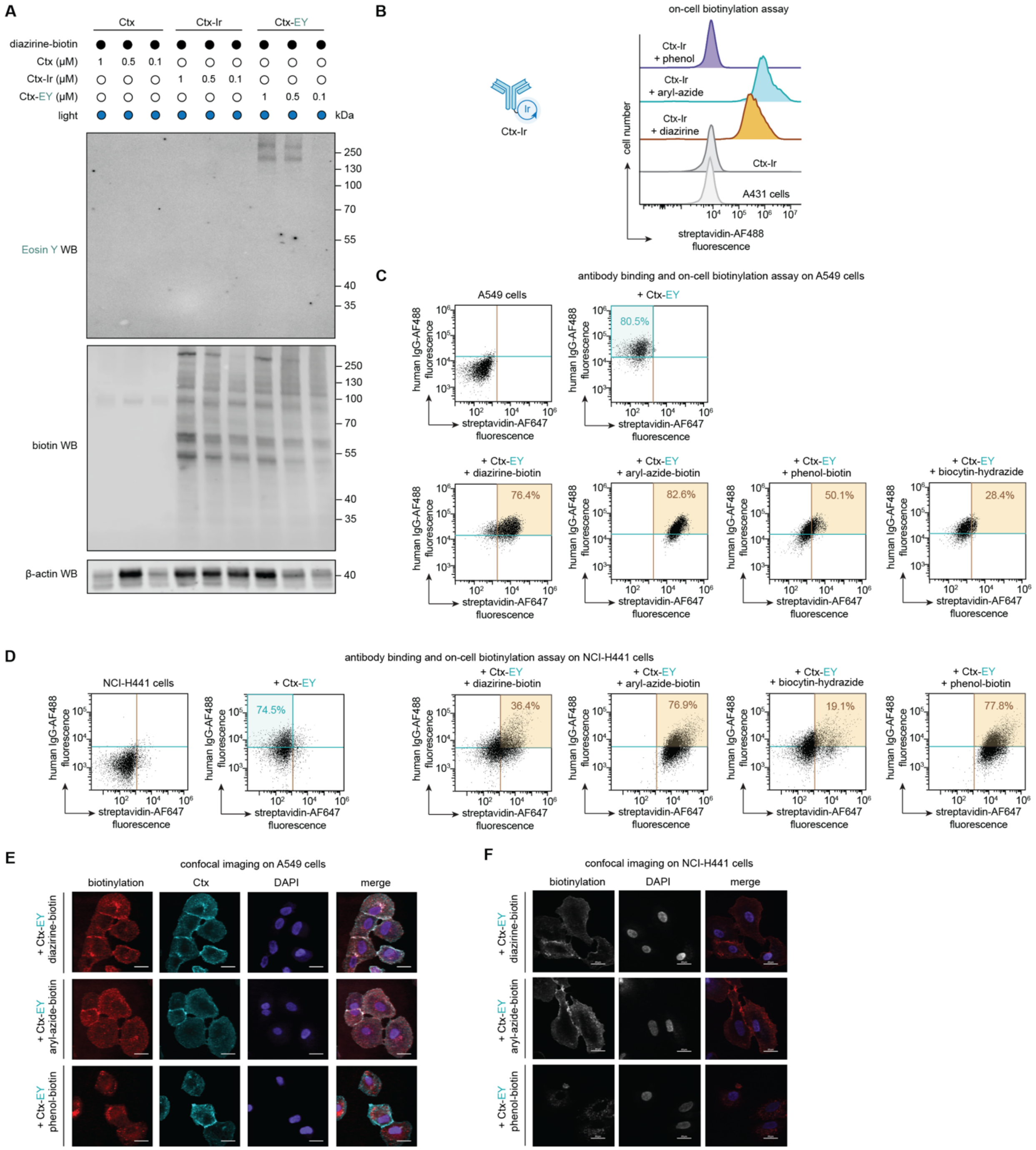


**Fig. S5. Ctx-EY can activate cell-surface biotinylation on A431, A549 and NCI-H441 cells using different photo-probes. (A)** On-cell biotinylation of A431 cells with diazirine-biotin from **Fig. 3D** confirmed by WB analysis. Dose-dependent labeling using Ctx-EY or Ctx-Ir are shown after 10 min blue LED illumination in the presence of 100 μM diazirine-biotin. **(B)** Flow cytometry analysis of on-cell biotinylation of A431 cells using Ctx-Ir confirmed that the iridium catalyst can activate biotin-diazirine and aryl-azide-biotin, but not phenol-biotin. **(C)** On-cell biotinylation with Ctx-EY on A549 cells expressing lower endogenous level of EGFR than A431 cells. **(D)** On-cell biotinylation with Ctx-EY on NCI-H441 cells expressing very low endogenous level of EGFR than either A549 or A431 cells. (**E)** Confocal microscopy imaging of cellular biotinylation and antibody binding with Ctx-EY on A549 cells expressing low endogenous level of EGFR. **(F)** Confocal microscopy imaging of cellular biotinylation with Ctx-EY on NCI-H441 cells expressing very low endogenous level of EGFR shows strong cell-surface labeling. Scale bar=20 μm. Immunoblot images are representative of at least two replicates.

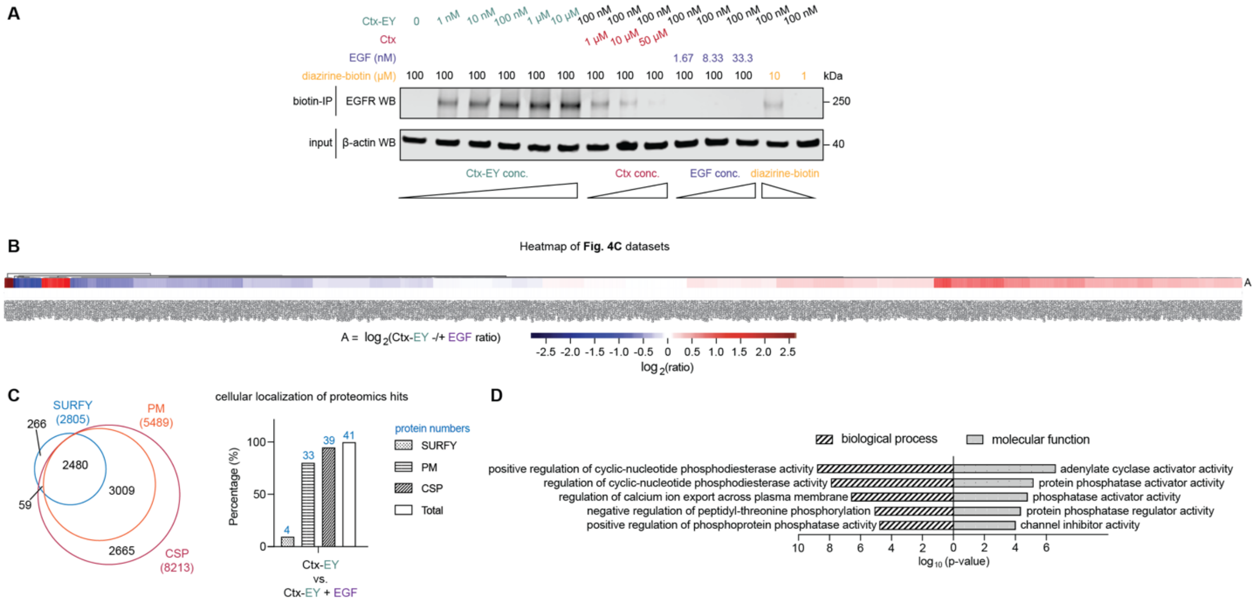


**Fig. S6. Specific labeling of EGFR labeling on A431 cells by Ctx-EY. (A)** Dose-dependent biotinylation of EGFR (shown by the wedges under each condition) in the presence of Ctx-EY, along with competition control conditions with either EGF or the unconjugated Ctx. Biotinylated proteins were enriched for each condition and analyzed by WB. Enrichment of EGFR is also dependent on the diazirine-biotin concentration. **(B)** Heatmap of the proteins identified in **Fig. 4C** by comparing enriched proteins using Ctx-EY versus Ctx-EY plus EGF competition. Significantly enriched proteins ($\log_2$(ratio)$\geq$1, p-value<0.05, unique peptide$\geq$2) are highlighted in red and listed in **Table S9**. **(C)** Cellular localization of protein hits discovered in **Fig. 4C**. All annotations were exported from UniProt. PM=plasma membrane, GO:0005886 (total protein number=5485, UniProt); CSP=cell surface proteome previously annotated (total protein number=8212) (*4*), SURFY=predicted surfaceome in silico as previously described (total protein number=2886) (*5*). **D)** Gene Ontology (GO) analysis performed on the EGFR-interacting candidates in **Fig. 4C**. Candidates were compared to a custom dataset containing UniProt-reviewed human proteome file with gene ontology localized at the plasma membrane (downloaded from UniProt database) using the methods previously described (*6*). Top 5 most significant biological process terms and molecular function terms were annotated with $\log_{10}$(p-value). Immunoblot images are representative of at least two biological replicates.

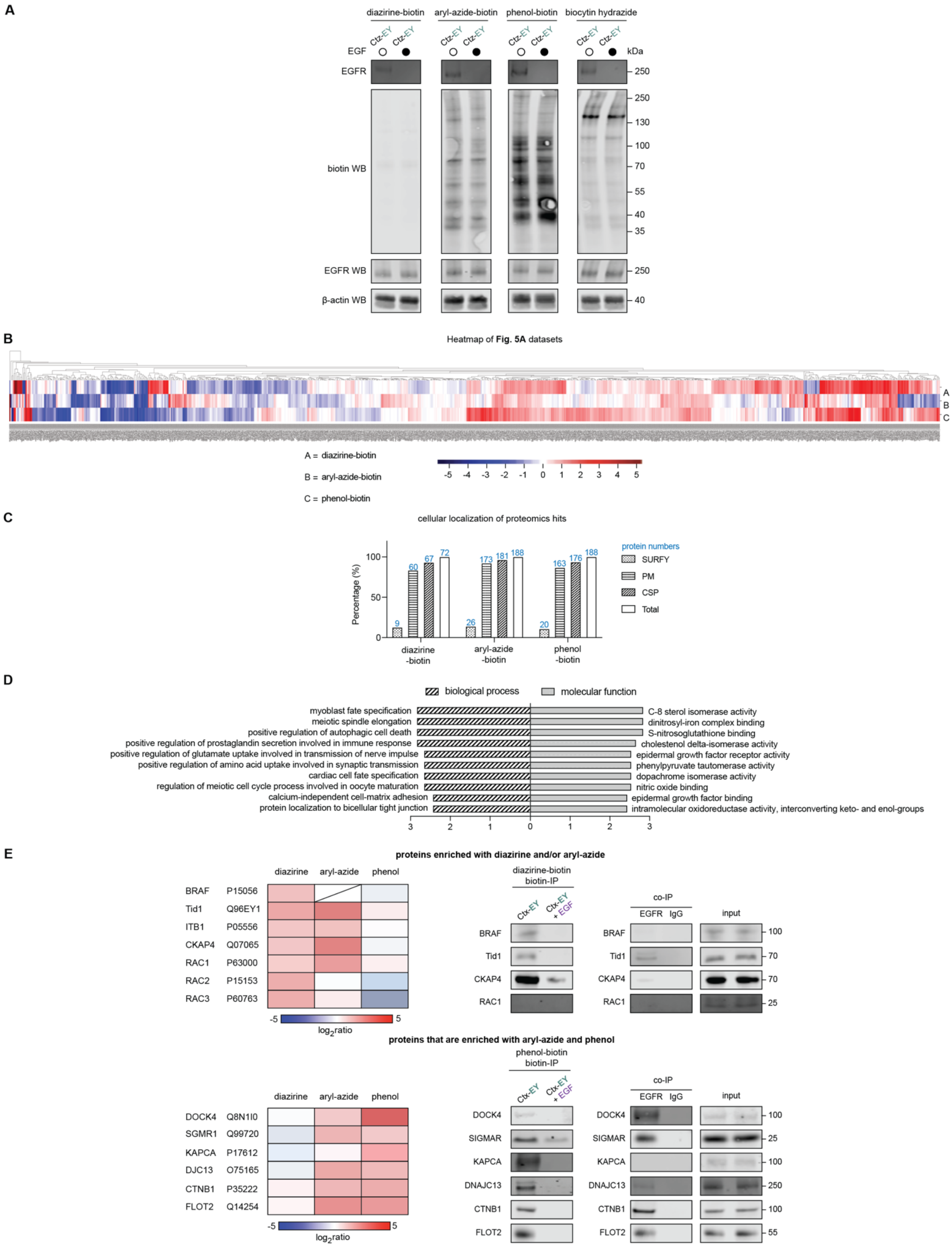


**Fig. S7. EGFR interactome profiling using Ctx-EY on live A549 cells. (A)** Full panel of WB analysis showing biotinylation of A549 cells using Ctx-EY showing it can activate biotinylation of A549 cells using biotin-diazirine, biotin-aryl-azide-, biotin-phenol and biocytin-hydrazide. **(B)** Heatmap of the EGFR-interacting candidates identified in **Fig. 5A**. Significantly enriched proteins ($log_2$(ratio)≥1, p-value<0.05, unique peptide≥2) are highlighted in red and listed in **Table S10-12**. **(C)** Categorized localization of protein hits discovered in **Fig. 5A**. Analysis was performed as described in **fig. S6C**. **(D)** Gene Ontology (GO) analysis performed on the EGFR-interacting candidates in **Fig. 5A**. Analysis was performed as described in **fig. S6D**.Top 10 most significant biological process terms and molecular function terms were annotated with p-value. **(E)** Enrichment ratios and validation of protein hits from the diazirine-biotin, aryl-azide-biotin or phenol-biotin dataset. All immunoblot images are representative of at least two biological independent experiments.

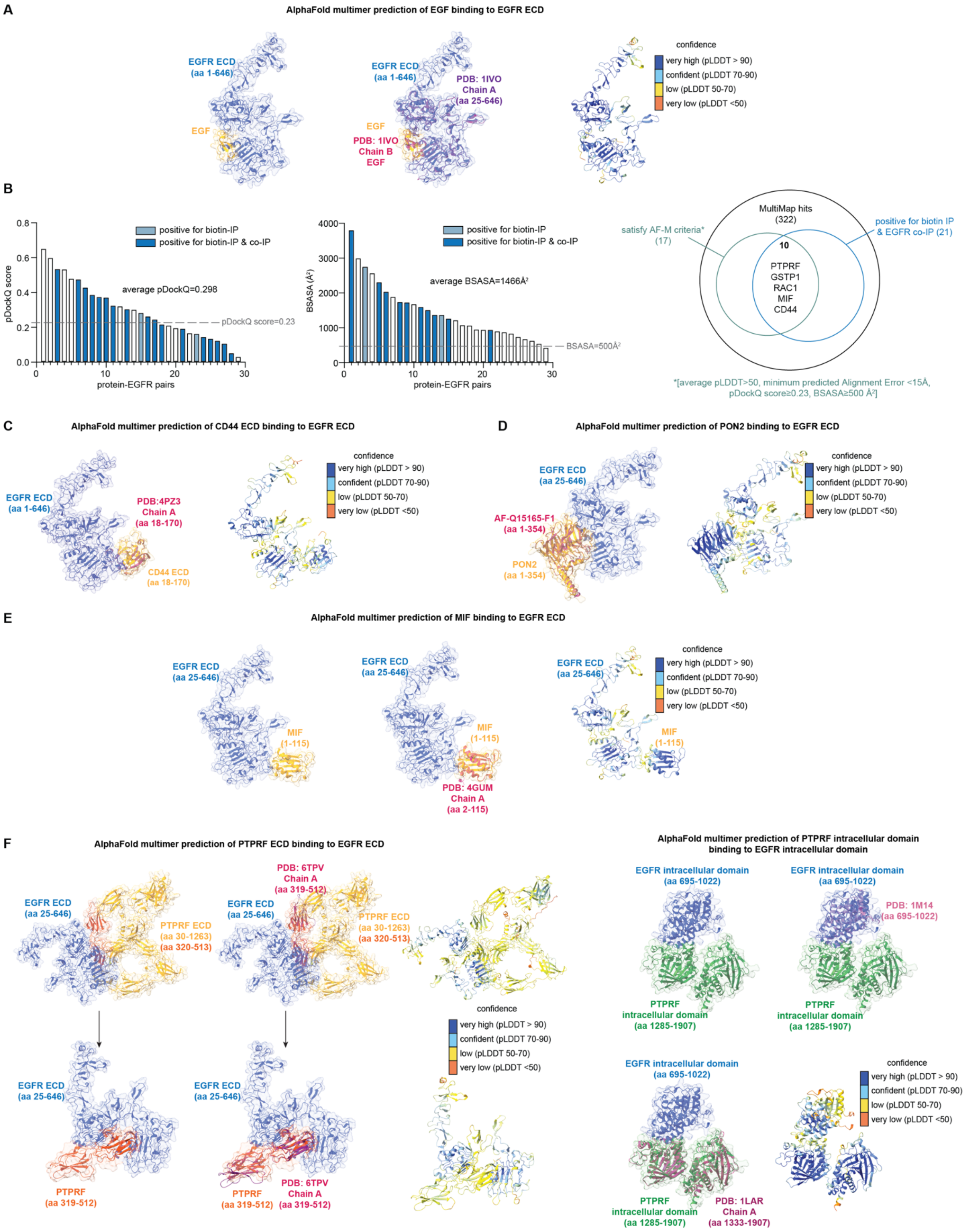


**Fig. S8. AlphaFold-Multimer predictions of EGFR binary complexes with EGFR interactors from MultiMap datasets. (A)** Predicted model of EGF bound to EGFR ECD via AlphaFold-Multimer compared with the crystal structure of EGF-bound EGFR (PDB: 1IVO, middle) and colored with pLDDT value (right). It is noteworthy that most residues were observed with high confidence (pLDDT>70). **(B)** Analysis of MultiMap hits using AlphaFold-Multimer and biochemical validation. An average pDockQ score of 0.298 and 1466Å$^2$ BSASA were observed for the 29 EGFR-protein pairs in the waterfall plots, both passing the criteria for highly confident heterodimeric AlphaFold-Multimer structures. **(C)** Predicted model of CD44 ECD bound to EGFR ECD via AlphaFold-Multimer shown in **Fig. 5F**. The predicted model was compared with the crystal structure of CD44 hyaluronan-binding domain in its ECD (PDB: 4PZ3, left) and colored with pLDDT value (right). **(D)** Predicted model of PON2 bound to EGFR ECD via AlphaFold-Multimer shown in **Fig. 5F**. Given that no crystal structures were available, the predicted structure was compared to the AlphaFold monomer prediction (AF-Q15165-F1, left) and colored with pLDDT value (right). **(E)** Predicted structure of MIF bound to the EGFR ECD via AlphaFold-Multimer. The predicted structure was compared with the crystal structure of MIF (PDB: 4GUM, middle) and colored with pLDDT value (right). **(F)** Predicted models of PTPRF ECD bound to EGFR ECD (left) and the PTPRF intracellular domain bound to the EGFR intracellular domain (right) via AlphaFold-Multimer shown in **Fig. 5F**. Predicted binding surface of PTPRF ECD (aa 319-512) was highlighted in orange-yellow and compared with the crystal structure of PTPRF ECD at the fibronectin type-III domain (PDB:6TPV). Both full PTPRF ECD and highlighted binding surface of PTPRF were colored with pLDDT value. Similarly, the predicted structure of the PTPRF intracellular domain bound to the EGFR intracellular domain was compared with the crystal structures of EGFR intracellular domain (PDB: 1M14) and PTPRF intracellular phosphatase domain (PDB: 1LAR), and colored with pLDDT value.

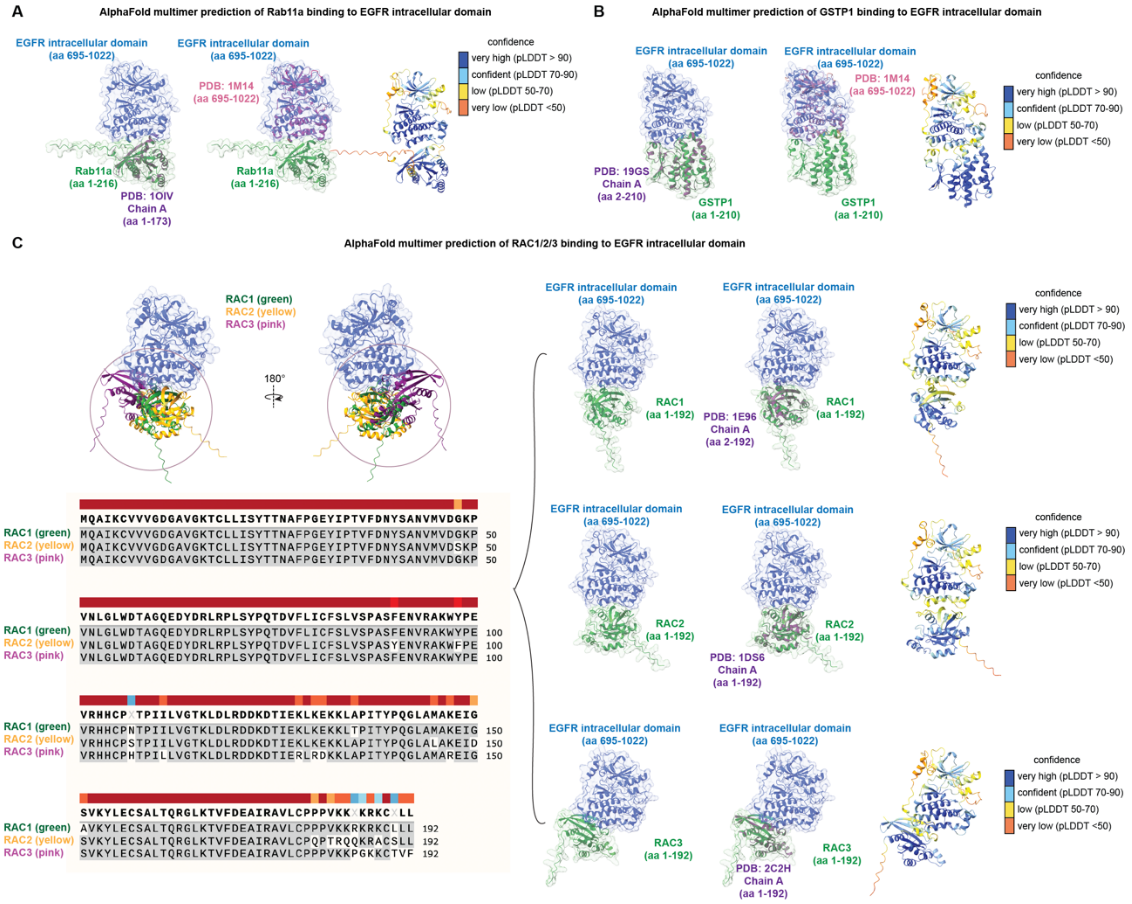


**Fig. S9. Additional predicted binary complexes of EGFR interactors with EGFR. (A)** Predicted model of Rab11a bound to the EGFR intracellular domain via AlphaFold-Multimer. The predicted structure was compared with the crystal structure of Rab11a (PDB: 1OIV, left), the crystal structure of the EGFR intracellular tyrosine kinase domain (PDB: 1M14, middle), and colored with pLDDT value (right). **(D)** Predicted model of GSTP1 bound to the EGFR intracellular domain via AlphaFold-Multimer shown in **Fig. 5F**. The predicted structure was compared with the crystal structure of GSTP1 (PDB: 19GS, left), the crystal structure of the EGFR intracellular domain (PDB: 1M14, middle), and colored with pLDDT value (right). **(C)** Predicted models of RAC1, RAC2 and RAC3 separately bound to the EGFR intracellular domain via AlphaFold-Multimer. Bound structures of the EGFR intracellular domain with RAC1 (top), RAC2 (middle), RAC3 (bottom) were compared with the crystal structures of RAC1 (PBD: 1E96), RAC2 (PDB: 1DS6), and RAC3 (PDB: 2C2H), and colored with pLDDT values. While sequence alignment showed high similarity (>88%) among three proteins, AlphaFold-Multimer provided distinct binding structures with the EGFR intracellular domain, which may explain why RAC1 was biotinylated with diazirine- and aryl-azide-biotin while RAC2 and RAC3 were only biotinylated with diazirine-biotin.

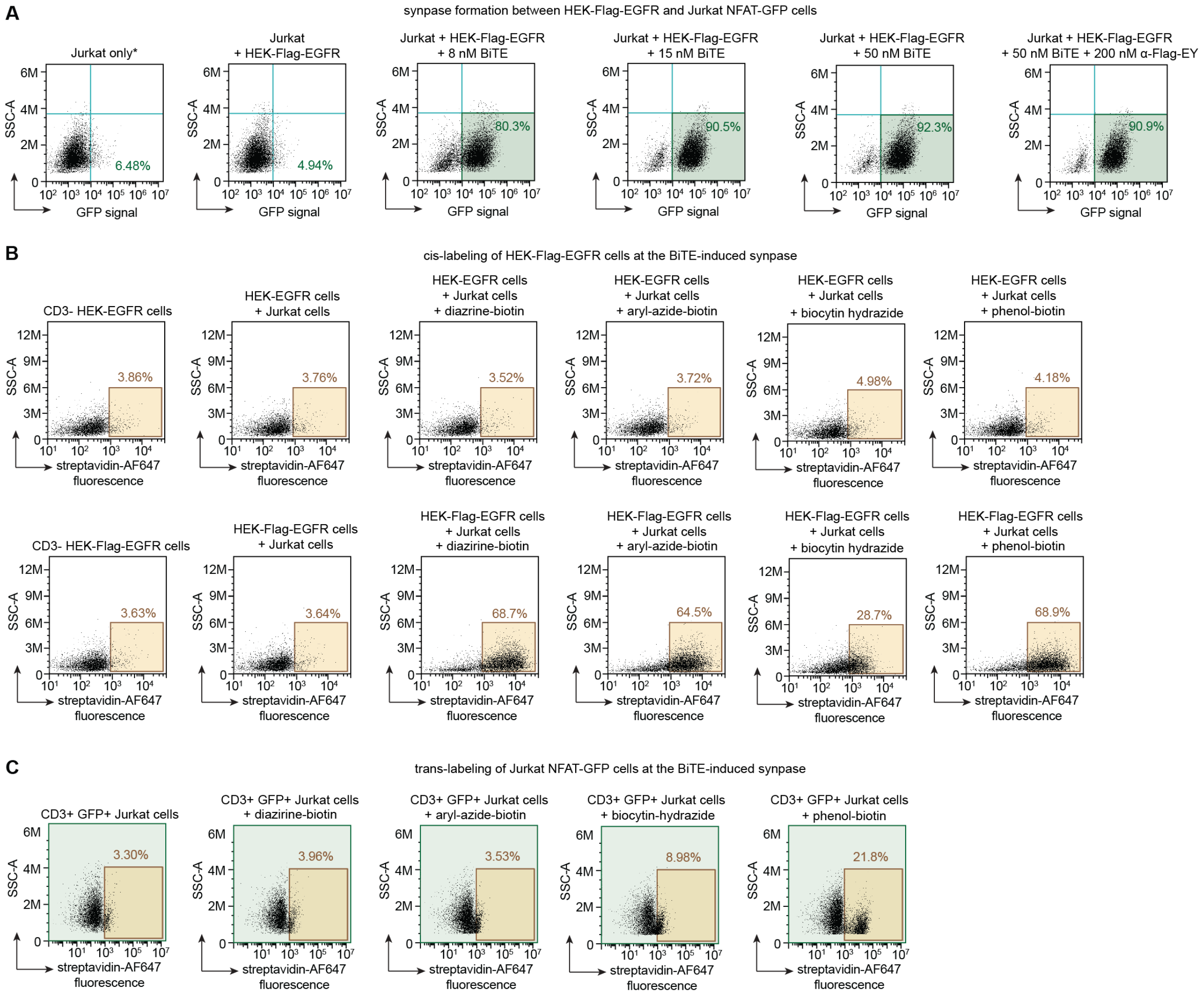


**Fig. S10. Targeted labeling of BiTE-induced synapses between Jurkat NFAT-GFP and HEK-Flag-EGFR using α-Flag-EY. (A)** Cell-cell engagement between Jurkat NFAT-GFP and HEK-Flag-EGFR induced by a bispecific T cell engager (BiTE) in a dose-dependent manner. Cell synapse formation was monitored by NFAT-GFP activation in Jurkat cells. Percentage of Jurkat cells engaging HEK293T-EGFR reached >90% and was not affected by the addition of EY-conjugated M1 Flag nanobody (α-Flag-EY). **(B)** Cis-labeling of HEK-Flag-EGFR using α-Flag-EY and four different photo-probes. HEK293T cells transfected with untagged EGFR (HEK-EGFR) and Flag-tagged EGFR (HEK-Flag-EGFR) were compared in parallel. Significant labeling with all four photo-probes in HEK-Flag-EGFR. **(C)** Trans-labeling of Jurkat cells using α-Flag-EY and four different photo-probes. CD3+ Jurkat cells were gated for GFP+ cells before the biotinylation levels were quantified. Only biotin-phenol provided significant trans-labeling shift.

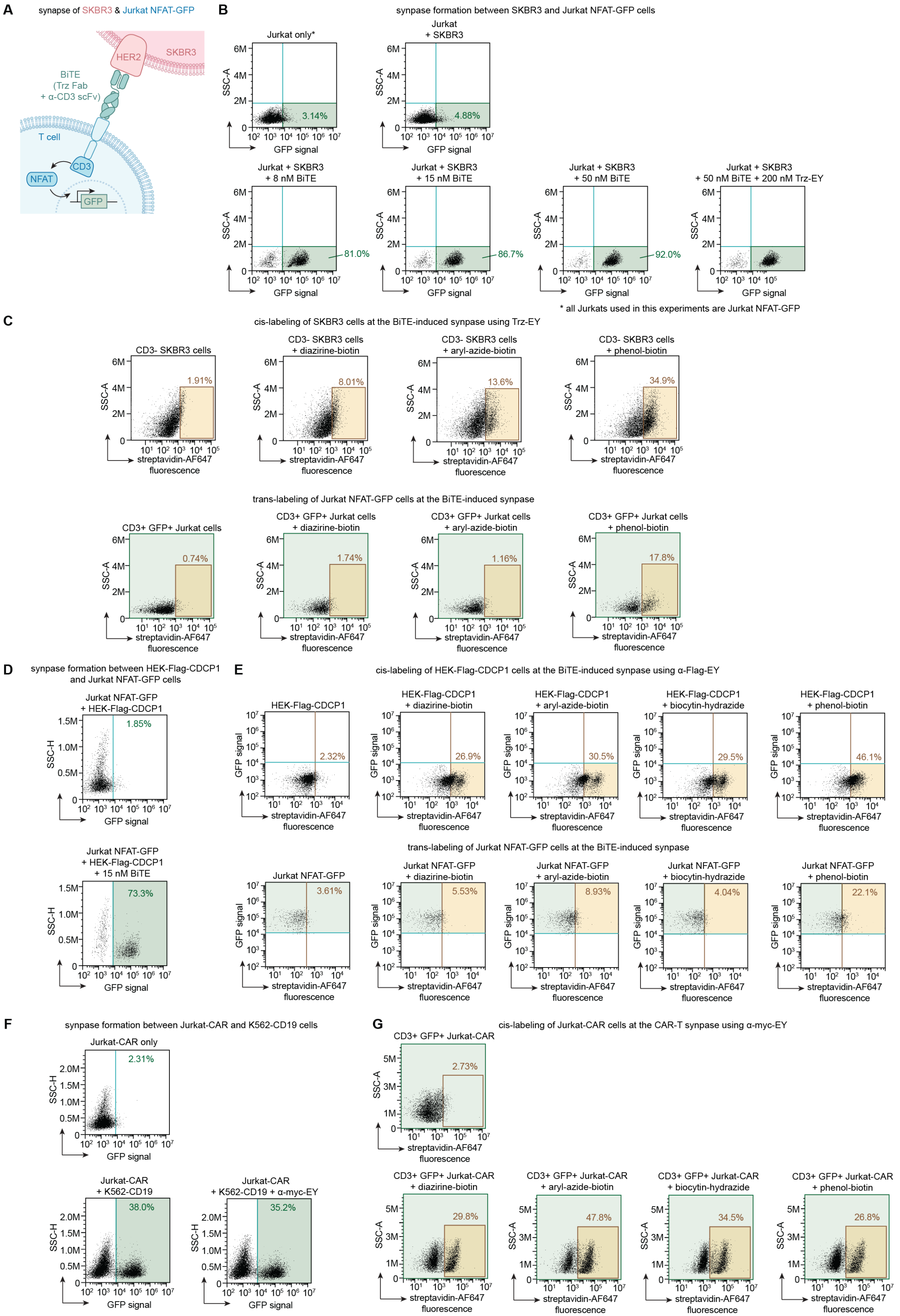
A
synapse of SKBR3 & Jurkat NFAT-GFP
HER2
SKBR3
BiTE
(Trz Fab
+ α-CD3 scFv)
T cell
CD3
NFAT
GFP
B
synapse formation between SKBR3 and Jurkat NFAT-GFP cells
Jurkat only*
3.14%
Jurkat
+ SKBR3
4.88%
Jurkat + SKBR3
+ 8 nM BiTE
81.0%
Jurkat + SKBR3
+ 15 nM BiTE
86.7%
Jurkat + SKBR3
+ 50 nM BiTE
92.0%
Jurkat + SKBR3
+ 50 nM BiTE + 200 nM Trz-EY
SSC-A
GFP signal
* all Jurkats used in this experiments are Jurkat NFAT-GFP
C
cis-labeling of SKBR3 cells at the BiTE-induced synapse using Trz-EY
CD3- SKBR3 cells
1.91%
CD3- SKBR3 cells
+ diazirine-biotin
8.01%
CD3- SKBR3 cells
+ aryl-azide-biotin
13.6%
CD3- SKBR3 cells
+ phenol-biotin
34.9%
streptavidin-AF647
fluorescence
trans-labeling of Jurkat NFAT-GFP cells at the BiTE-induced synapse
CD3+ GFP+ Jurkat cells
0.74%
CD3+ GFP+ Jurkat cells
+ diazirine-biotin
1.74%
CD3+ GFP+ Jurkat cells
+ aryl-azide-biotin
1.16%
CD3+ GFP+ Jurkat cells
+ phenol-biotin
17.8%
D
synapse formation between HEK-Flag-CDCP1
and Jurkat NFAT-GFP cells
Jurkat NFAT-GFP
+ HEK-Flag-CDCP1
1.85%
Jurkat NFAT-GFP
+ HEK-Flag-CDCP1
+ 15 nM BiTE
73.3%
SSC-H
GFP signal
E
cis-labeling of HEK-Flag-CDCP1 cells at the BiTE-induced synapse using α-Flag-EY
HEK-Flag-CDCP1
2.32%
HEK-Flag-CDCP1
+ diazirine-biotin
26.9%
HEK-Flag-CDCP1
+ aryl-azide-biotin
30.5%
HEK-Flag-CDCP1
+ biocytin-hydrazide
29.5%
HEK-Flag-CDCP1
+ phenol-biotin
46.1%
trans-labeling of Jurkat NFAT-GFP cells at the BiTE-induced synapse
Jurkat NFAT-GFP
3.61%
Jurkat NFAT-GFP
+ diazirine-biotin
5.53%
Jurkat NFAT-GFP
+ aryl-azide-biotin
8.93%
Jurkat NFAT-GFP
+ biocytin-hydrazide
4.04%
Jurkat NFAT-GFP
+ phenol-biotin
22.1%
F
synapse formation between Jurkat-CAR and K562-CD19 cells
Jurkat-CAR only
2.31%
Jurkat-CAR
+ K562-CD19
38.0%
Jurkat-CAR
+ K562-CD19 + α-myc-EY
35.2%
G
cis-labeling of Jurkat-CAR cells at the CAR-T synapse using α-myc-EY
CD3+ GFP+ Jurkat-CAR
2.73%
CD3+ GFP+ Jurkat-CAR
+ diazirine-biotin
29.8%
CD3+ GFP+ Jurkat-CAR
+ aryl-azide-biotin
47.8%
CD3+ GFP+ Jurkat-CAR
+ biocytin-hydrazide
34.5%
CD3+ GFP+ Jurkat-CAR
+ phenol-biotin
26.8%

**Fig. S11. Extended applications of targeted cell synapse labeling. (A)** Scheme of on-cell labeling of Jurkat NFAT-GFP and SKBR3 induced by a BiTE that recognizes HER2. **(B)** Cell-cell engagement between Jurkat NFAT-GFP and SKBR3 cells was monitored by NFAT-GFP activation. Percentage of Jurkat cells engaging SKBR3 reached >90% and the synapse was not affected by the addition of the EY-conjugated Trz (Trz-EY). **(C)** Cis- and trans-labeling SKBR3 and Jurkat NFAT-GFP between a BiTE-induced synapse using α-HER2-EY and three different photo-probes. **(D)** Cell-cell engagement between HEK-Flag-CDCP1 and Jurkat NFAT-GFP monitored by NFAT-GFP activation. **(E)** Cis- and trans-labeling of HEK-Flag-CDCP1 and Jurkat NFAT-GFP between a BiTE-induced synapse using α-Flag-EY. **(F)** Cell-cell engagement between Jurkat-CAR with K562-CD19 monitored by NFAT-GFP activation. **(G)** Cis-labeling of Jurkat-CAR with K562-CD19 shown in **Fig. 6D**.

**Methods**

**General methods and instrumentation.**

Illumination was performed using a PennOC Photoreactor M1 for indicated time using a 450 nm blue light source at 100% intensity, or Thor Labs LED Array light source (LIU470A for 470 nm LED array, LIU525B for 525 nm LED array) along with a LED mounting adapter (AD38) for indicated time. Flow cytometry experiments were performed on a CytoFlex flow cytometer (Beckman CytoFlex) and analyzed using FlowJo software. Cell sorting experiments were performed on a SONY SH800 cell sorter. Proteomics experiments were performed on a TimsTOF PRO (Bruker) equipped with a CaptiveSpray source and a nanoElute System. The peptides were separated on a 25 cm, ReproSil c18 1.5 µM 100 Å column (PepSep, PN. # PSC- 25-150-15-UHP-nc). All acquired data was searched using PEAKS online Xpro 1.6 (Bioinformatics Solutions Inc). Protein quantification was performed by bicinchoninic acid assay on a multimode microplate reader Infinite 200 PRO (Tecan Trading AG, Switzerland). Sonication of cells or protein pellets was performed using a QSonica Q500 Sonicator (QSonica Sonicators, Newtown, CT). DNA, RNA or protein concentrations were measured using a NanoDrop 2000 spectrophotometer (Thermo Scientific).

For immunoblotting analysis, proteins were loaded on 4-12% BisTris gels (Bolt 4-12% 17-well, Thermo Fischer, NW04127BOX), and transferred from SDS-PAGE gels to PVDF membranes (Thermo Fischer, IB24002) using an iBlot-2 dry blotting system (Thermo Scientific, IB21001). Membranes were blocked with Tris buffered saline (TBST, 37mM sodium chloride, 20mM Tris, 2.7mM potassium chloride, 0.05% Tween 20; pH=7.4) containing 0.1% Tween-20 and 5% BSA and incubated with the primary antibodies and the secondary antibodies sequentially including anti-rabbit IgG Goat IR800 secondary antibody (Rockland, 926-32211), anti-rabbit IgG Goat IR680 secondary antibody (Rockland, 611-144-002), anti-rabbit IgG Goat secondary antibody peroxidase (Rockland, 611-1302), anti-mouse IgG Goat IR800 secondary antibody (Rockland, 610-145-211) and anti-mouse IgG Goat IR680 secondary antibody (Rockland, 610-144-002). Immunoblots images were captured by an infrared LI-COR imager (Odyssey CLx). In-gel fluorescence and immunoblot fluorescence signals were detected on a BioRad imager (ChemiDoc XRS+ System).

**General chemical methods and instrumentation.**

Chemicals were purchased including TFPA-$PEG_3$-biotin (Thermo Scientific, 21303), biotinyl tyramide (Sigma-Aldrich, SML2135), eosin-5-isothiocyanate (Biotium, 90091), 5-iodoacetamidoerythrosin (Alfa Chemistry, ALP3853), erythrosine B disodium salt (Alfa Aesar, A14180-14), DBCO-$PEG_4$-amine (Click Chemistry Tools, A103P-100; separately synthesized by ChemPartner), 2-(Prop-2-yn-1-yloxy)-4-(3-(trifluoromethyl)-3H-diazirin-3-yl)benzoic acid (Sigma-Aldrich, 900858), Eosin Y (Sigma-Aldrich, E4009) and rose Bengal (Sigma-Aldrich, 33000). All solvents and reagents were purchased from chemical suppliers (Sigma Aldrich, Acros Organics, Thermo Scientific or VWR Chemicals BDH®) and were used as received unless otherwise noted. Flash Column Chromatography was performed using Teledyne ISCO CombiFlash EZ Prep chromatography system, employing pre-packed silica gel Teledyne ISCO RediSep cartridges. Protein mass spectra were obtained using a Waters Xevo G2-XS time-of-flight mass spectrometer operating with Waters MassLynx software (version 4.2). DBCO-$PEG_3$-EY was synthesized in house before it was mass produced and characterized by ChemPartner. Diazirine-biotin (diazirine-$PEG_3$-biotin) was synthesized and characterized by Medicilon according to literature (*7*).

Proton nuclear magnetic resonance spectrum ($^1H$ NMR) and carbon nuclear magnetic resonance spectrum ($^{13}C$ NMR) were recorded on a Bruker 400 MHz instrument at 25 °C. Chemical shifts were reported in parts per million (ppm, δ scale) relative to residual solvent as an internal reference (DMSO: 2.50 ppm for $^1H$ and 39.52 ppm for $^{13}C$). Data are represented as follows: chemical shift, multiplicity (s = singlet, d = doublet, t = triplet, q = quartet, quin = quintet, m = multiplet and/or multiple resonances, br = broad, app = apparent), integration, coupling constant (*J*) in Hertz (Hz), and assignment. Infrared (IR) spectrum was recorded on a Bruker ALPHA FT-IR and are reported in terms of frequency of absorption ($cm^{-1}$) and intensity of absorption (s = strong, m = medium, w = weak, br = broad).

**Reaction monitoring by LC-MS.**
Reaction catalyzed by EY was monitored and quantified via LC-MS. In general, 100 µL reaction systems in water with 1 µM EY and 10 µM 2-(Prop-2-yn-1-yloxy)-4-(3-(trifluoromethyl)-3H-diazirin-3-yl)benzoic acid, or EZ-Link™ TFPA-$PEG_3$-Biotin (Thermo Scientific, 21303) were mixed in an Eppendorf tube. The reactions were illuminated with indicated LED at 4 °C and monitored over time. At the indicated time points, 1 µL of the reaction product was diluted with 100 µL with an acetonitrile:water mixture (v/v 50%) and 1 µL of the final mixture was injected onto a Water Acquity UPLC BEH C18 1.7 µm column and eluted with a linear gradient of 5-95% acetonitrile/water (with 0.1% formic acid) over 3 min. Chromatograms were recorded with a time-of-flight mass spectrometer (Waters Xevo G2-XS).

**Antibodies and biological reagents.**
Antibodies were purchased including: Ctx (cetuximab, Selleck Chemicals, A2000), Trz (trastuzumab, Selleck Chemicals, A2007), anti-EGFR (Thermo Scientific, MA5-13319; Cell Signaling Technology, 4267S), anti-HER2 (Cell Signaling Technology, 2165S), anti-MIF (Proteintech, 20415-1-AP), anti-GSTP1 (Proteintech, 15902-1-AP), anti-ZO1 (Proteintech, 21772-1-AP), anti-PGRMC1 (Cell Signaling Technology, 13856T), anti-Integrin β1 (Cell Signaling Technology, 4706S), anti-β-actin (Santa Cruz Biotechnology, sc-47778), anti-LGALS3 (Cell Signaling Technology, 12733S), anti-CD44 (Cell Signaling Technology, 3578S), anti-ADDB (Proteintech, 14640-1-AP), anti-PON2 (Abcam, ab183710), anti-PTPRF (anti-LAR, R&D system, MAB3004-SP), anti-Rab11a (Cell Signaling Technology, 2413S), anti-RHOC (Cell Signaling Technology, 3430T), anti-PDCD6IP (Proteintech, 12422-1-AP), anti-BRAF (Cell Signaling Technology, 14814S), anti-Tid1 (Cell Signaling Technology, 4775S), anti-CKAP4 (Proteintech, 16686-1-AP), anti-Rac1/2/3 (Cell Signaling Technology, 2465T), anti-DOCK4 (Proteintech, 21861-1-AP), anti-SIGMAR1 (Proteintech, 15168-1-AP), anti-PKACa (Cell Signaling Technology, 4782S), anti-DNAJC13 (Bethyl Laboratories, A304-872A), anti-β-catenin (Cell Signaling Technology, 8480T), anti-Flot2 (anti-Flotilin 2, Cell Signaling Technology, 3436S), mouse IgG1 isotype control (BD Biosciences, 556648), anti-CD3D (Cell Signaling Technology, 31857S), anti-CD19 (Cell Signaling Technology, 90176T), anti-CDCP1 (Cell Signaling Technology, 4115S) and anti-Myc (Santa Cruz Biotechnology, sc-40). Anti-M1-FLAG antibody was purified in HEK293T cells using the sequence gifted by the Kruse lab (Harvard Medical School).

The following antibodies were used in flow cytometry assays: anti-CD3-AlexaFluor561 (Thermo Scientific, 505-0038-41), anti-CD3-PE (BioLegend, 300456), anti-CD19-PE (BioLegend, 302254), streptavidin-AlexaFluor488 (Thermo Scientific, S32354), streptavidin-AlexaFluor647 (BioLegend, 405237), anti-EGFR-AlexaFluor647 (Fisher Scientific, 352918), anti-human IgG-AlexaFluor488 (BioTechne, FAB110G) and anti-human IgG-AlexaFluor647 (BioTechne, FAB110R). Recombinant proteins included human EGFR (Bio-Techne, 1095-ER-002) and human HER2 (Acro Biosystems, HE2-H5225). Gels were imaged with InstantBlue protein stain (Expedeon, ISB1L). Albumin was purchased from Sigma-Aldrich (A1887). For enrichment assays, NeutrAvidin agarose beads (Pierce, 29200) and protein A magnetic beads (Cell Signaling Technology, 73778) were used. Cell lysis buffer was prepared by diluting from 10X cell lysis buffer (Cell Signaling Technology, 9803S) or from 10X RIPA buffer (EMD Millipore, 20-188). Sample loading buffer were diluted from 4X LDS sample loading buffer (G Biosciences, 786-323). Sequencing-grade modified trypsin (Promega, V5111), sequencing-grade chymotrypsin (Promega, V1061) and mini Bio-Spin columns (Bio-Rad, 7326207) were purchased. When performing solvent exchange processes, 7 kDa Zeba Spin desalting columns (Thermo Fischer, 89883) were used.

**Plasmid Construction.**
Plasmids for the Ctx-OKT3 BiTE, Trz-OKT3 BiTE and α-CDCP1-OKT3 BiTE that targeted EGFR, HER2 and CDCP1, respectively were constructed by standard molecular biology methods and as previously described (*8*). For example, DNA fragments of Ctx Fab heavy and light chain were synthesized by integrated DNA technologies (IDT). OKT3 scFv was amplified using primers shown below. All BiTEs were constructed in the pFUSE-hlgG1 vector (InvivoGen) with IL-2 signal peptide for mammalian expression. Ctx Fab heavy chain was cloned on one vector, and the

Fab light chain genetically fused with the N-terminus of OKT3 was cloned on a separate copy of the vector. The sequence of the linker between the light chain and scFv is as follows: GGGGS. All sequences were confirmed by Sanger (Quintarabio) and whole-plasmid (Primordium Labs) sequencing.
Cloning primers:
(forward) 5'-CCGGGGGGAATGTGGCGGCGGAGGCAGCGACATCAAGCTGCAGCA-3'
(reverse) 5'-ATCTTATCATGTCTGGCCAGCTAGCTCACTTCAGTTCCAGCTTTG-3'

**Cell culture.**
A549, A431, NCI-H441, SKBR3, Jurkat and HEK293T cells were all purchased from the UCSF cell culture facility. A549, A431, NCI-H441 and SKBR3 cells were cultured and maintained in ATCC recommended conditions. HEK293T cells with Flag-CDCP1 overexpressed were generated according to literature (*8*). HEK293T cells with Flag-EGFR overexpressed were generated similarly. Jurkat cells expressing NFAT-GFP reporter were cultured in RPMI containing 10% FBS, 1% pen/strep and 2 mg/mL geneticin. K562-CD19 and Jurkat-CAR cells were cultured according to literature (*9*).

**Mammalian protein expression.**
HEK293Expi (Expi293) cells were cultured in FreeStyle Expi293 media (Gibco, 12338018) at 37 ˚C and 8% humidity with orbital shaking at 250 rpm. Protein expression plasmids were cloned into a pFUSE vector (InvivoGen) with upstream IL-2 secretion signal. Cells were transfected at 3M/mL density using FectoPRO transfection kit (Genesee Scientific, 55-332) according to manufacturers' instructions. After expression for 4-6 days, the supernatant from Expi293 cells was collected by centrifuging at 4000 g for 30 min and filtered through a 0.45 μm filter. After equilibrating Hitrap Protein A/L affinity column (GE Healthcare, 12-0402-01) or nickel resin, columns were washing with PBS (pH 7.4) using six times the column volume, and protein was eluted into 100 mM acetic acid. Following pH neutralization, the purified proteins were buffer exchanged with PBS (pH 7.4) using 10 kDa MW spin filters (AmiconUltra, UFC9010). Protein samples prepared in 4X loading dye with or without DTT were then characterized using SDS-PAGE. Purified proteins were quantified using A280 channel on a NanoDrop, and flash frozen in single use aliquots for storing at -80 ˚C or used fresh within a week.

**General protocol for antibody conjugation with EY.**
In order to generate antibody-EY conjugates, different bioconjugation strategies were employed in **Fig. 2** including NHS labeling and oxaziridine labeling. For NHS labeling, a 200 μL reaction mixture was prepared with final concentrations of 10 μM purified antibody and 50 μM N-Hydroxysuccinimidyl-4-azidobenzoate (NHS-azide, Santa Cruz Biotechnology, sc-263835) along with 10 mM sodium bicarbonate in PBS. The reaction was incubated for 1 h at 25 ˚C before another portion of NHS-azide was added to reach a final concentration 100 μM. The resulting mixture was allowed to react for additional 1 h at 25 ˚C. Then the conjugate was purified using a 7 kDa Zeba Spin desalting column (Thermo Scientific, 89882). The resulting azide-conjugated antibodies were then incubated with 100 μM DBCO-PEG$_4$-EY for 16 h at 4 ˚C before purification with a 7 kDa Zeba Spin desalting column twice. Formation of the desired antibody-NHS-EY conjugates was confirmed by LC-MS, SDS-PAGE and UV-Vis spectrum scanning. The concentrations of proteins were calculated from SDS-PAGE gels. After characterization, the antibody-NHS-EY conjugate was flash frozen for future usage or used fresh within a week. For oxaziridine labeling, a 200 μL

reaction was prepared with 10 μM purified antibody and 50 μM oxaziridine-azide (piperidine-oxaziridine 8 synthesized accordingly literature) (*3*) in PBS. The reaction was incubated for 1 h at 25 ˚C before purification using a 7 kDa Zeba Spin desalting column. The resulting azide-conjugated antibodies were then incubated with 50 μM DBCO-PEG$_4$-EY for 16 h at 4 ˚C before purification with a 7 kDa Zeba Spin desalting column twice. Formation of the desired antibody conjugate was confirmed as described above. After characterization, the corresponding antibody-Ox-EY conjugate was flash frozen for future usage or used fresh within a week.

**Western blot protocol.**
Cells were incubated at 37 °C in 5% $CO_2$ to 80% confluency and washed with 5 mL PBS three times before they were incubated with PBS with 0.04% EDTA (free of calcium and magnesium) for 15 min. Dissociated cells were collected and washed with 10 mL PBS three times before they were pelleted via centrifugation at 300 g for 5 min in 1.5 mL Eppendorf tubes. Cell pellets were resuspended in 1 mL 1X RIPA lysis buffer (EMD Millipore) supplemented with 1X cOmplete™ protease inhibitor cocktail (Roche). After 15 min incubation on ice, cells were sonicated for 15 sec (5 sec on, 5 sec off, 20%). Cell lysates were then cleared by centrifugation at 20,000 g for 10 min at 4 ˚C. Protein concentrations were measured using a BCA assay kit (Pierce). Samples were then analyzed by SDS-PAGE and transferred onto PVDF membranes using an iBlot2 transfer stack. Total protein was first assessed using Ponceau S staining. The membranes were then blocked using TBST with 5% BSA for 1 h at 25 ˚C before primary and secondary antibodies were added. Biotinylation and near-infrared Western blot imaging were conducted using an Odyssey Li-COR imaging system before further analysis using ImageStudioLite.

**General flow cytometry.**
Cultured cells or co-culture systems were incubated at 37 °C in 5% $CO_2$ for the duration of the assay. Cells were first washed three times with 5 mL PBS followed by additional three washes with 5 mL filtered 3% BSA in PBS. Then the cells were either directly resuspended in 0.5 mL PBS for flow cytometry analysis or stained with corresponding primary antibody for 1 h at 4 °C. The stained cells were then washed three times with 5 mL filtered 3% BSA in PBS before they were resuspended in 0.5 mL PBS for flow cytometry analysis. Flow cytometry data were analyzed on FlowJo.

**On-cell antibody binding and biotinylation assay.**
A431, A549 or NCI-H441 cells were incubated at 37 °C in 5% $CO_2$ to 80% confluency and washed with PBS three times before they were incubated with PBS with 0.04% EDTA (free of calcium, magnesium) for 15 min. Dissociated cells were collected and washed three times with 10 mL PBS before they were pelleted in 1.5 mL Eppendorf tubes. Cells were resuspended in PBS to 1 X $10^6$ cell/mL concentration, and then incubated with or without antibody or reagents as indicated at 4 °C. Mixtures were illuminated with LED, pelleted and washed again three times with 0.5 mL PBS. Treated cells were then stained with streptavidin-AlexaFluor488 (1:2000 diluted with filtered 3% BSA in PBS) and/or anti-human IgG-AlexaFluor647 (1:2000 diluted with filtered 3% BSA in PBS) for 1 h at 4 °C before they were washed three times with 0.5 mL filtered 3% BSA in PBS. Samples were then suspended in 0.5 mL PBS before flow cytometry analysis.

**Recombinant protein biotinylation assay.**

In order to test the catalytic function of EY when conjugated on an antibody, both self-labeling and target-biotinylation were validated using recombinant proteins. A 100 μL reaction system in PBS was prepared with 10 μM purified antibody or antibody-EY conjugate with or without equivalent amount of binding antigen for 15 min at 4 °C. For example, 10 μM Ctx or Ctx-EY conjugate was incubated with or without 10 μM recombinant EGFR in PBS. Photo-probe (diazirine-biotin, aryl-azide-biotin, biocytin-hydrazide, or biotin-phenol) was then added into the solution to reach a final concentration of 100 μM and mixed thoroughly before illumination with LED for 10 min at 4 °C. Afterwards, proteins were precipitated with pre-chilled acetone to get rid of excess small molecules, resuspended in PBS or sample loading buffer, and subjected to SDS-PAGE or LC-MS/MS sample preparation.

**General protocol for antibody-EY labeling on cells.**
A431, A549 or NCI-H441 cells were incubated at 37 °C in 5% CO2 to 80% confluency and washed three times with PBS before they were incubated with PBS with 0.04% EDTA (free of calcium, magnesium) for 15 min. Dissociated cells were collected and washed with 5 mL PBS three times before they were pelleted in 1.5 mL Eppendorf tubes and resuspended in pre-chilled PBS to 10M cell/mL concentration. Indicated amounts of antibody-NHS-EY conjugates were pre-chilled and added to the cells for 15 min at 4 °C before excessive antibody-EY conjugates were removed by washing with 1 mL pre-chilled PBS. The antibody-bound cells were then resuspended in 1 mL pre-chilled PBS. Photo-probe (diazirine-biotin, aryl-azide-biotin, biocytin-hydrazide, or biotin-phenol) was then added into the cell solution to reach a final concentration of 100 μM and mixed thoroughly before illumination with LED for 10 min at 4 °C. Afterwards, cells were pelleted again and subjected to flow cytometry or LC-MS/MS sample preparation.

**Sample preparation for LC-MS/MS analysis.**
For sample processing, cell pellets were resuspended in 1 mL 1X RIPA lysis buffer (EMD Millipore) supplemented with 1X cOmplete™ protease inhibitor cocktail (Roche). After 15 min incubation on ice, cells were sonicated for 15 sec (5 sec on, 5 sec off, 20%). Cell lysates were then cleared by centrifugation at 20,000 g for 10 min at 4 ˚C. Protein concentrations in the cleared supernatant were measured using a BCA assay kit (Pierce). Proteins were then added to 200 μL NeutrAvidin agarose beads (Pierce) that were pre-washed with 5 mL PBS for 3 times and incubated for 16 h at 4 ˚C. Afterwards, supernatant was discarded using mini Bio-spin columns (Bio-Rad) and the beads were washed three times with 3 mL 1X RIPA lysis buffer, three times with 3 mL 1X PBS with 1M NaCl, and three time with 3 mL of freshly prepared 2M urea in 50 mM ammonium bicarbonate. The beads were then suspended in 100 μL PBS to re-constitute 50% slurry with 10 μL bead slurry separated for Western blotting.

Proteins on the washed beads were then digested using the Preomics iST kit in an on-bead digestion format according to the manufacturer's instructions. In brief, washed beads were suspended in 100 μL LYSE buffer provided by Preomics iST kit and incubated at 55 ˚C for 10 min for reduction and alkylation. Once the beads cooled down to room temperature, 50 μL of pre-reconstituted DIGEST were added to the beads and incubated at 37 ˚C for 3 h with shaking. The digested peptides were then collected using mini Bio-Spin columns (Bio-Rad) and another 50 μL of LYSE buffer were added to wash the beads. Afterwards, 100 μL of STOP solution was added to the combined flow-through elution and mixed using vigorous vortexing. Then the peptides were desalted using the Preomics desalting columns before they were dried under vacuum and resuspended in 15 μL solvent A (0.1% formic acid with 2% acetonitrile) for mass spectrometry analysis. Peptide amount was monitored by quantitative fluorometric peptide assay (Pierce).

**Proteomics analysis of digested peptide samples.**
Proteomics experiments were performed on a TimsTOF PRO (Bruker) equipped with a CaptiveSpray source and a nanoElute system. The peptides were separated on a 25 cm, ReproSil c18 1.5 µM 100 Å column (PepSep, PN. # PSC-25-150-15-UHP-nc) using a step-wise linear gradient method with water in 0.1% formic acid (solvent A) and acetonitrile with 0.1% formic acid (solvent B): 5-30% solvent B for 90 min at 0.5 µl/min, 30-35% solvent B for 10 min at 0.6 µl/min, 35-95% solvent B for 4 min at 0.5 µl/min, 95% hold for 4 min at 0.5 µl/min). Acquired data was collected in a data-dependent acquisition mode with ion mobility activated in PASEF mode. MS and MS/MS spectra were collected with m/z ranging from 100 to 1700 in positive mode.

**Analysis of proteomics dataset.**
All acquired data was searched using PEAKS online Xpro 1.6 (Bioinformatics Solutions Inc.). Spectral searches were performed using a custom FASTA-formatted dataset containing Swiss Uniprot-reviewed human proteome file with gene ontology localized the plasma membrane (downloaded from UniProt database). A precursor mass error tolerance was set to 20 ppm and a fragment mass error tolerance was set at 0.03 ppm. Peptides, ranging from 6 to 45 amino acids in length, were searched in semi-specific trypsin digest mode with a maximum of three missed cleavages. Carbamidomethylation (+57.0214 Da) on cysteines was set as a static modification while methionine oxidation (+15.9949 Da) and lysine acetylation (+42.0115 Da) were set as a variable modification. Peptides were filtered based on a false discovery rate (FDR) of 1%. Samples were normalized using total ion current (TIC). Protein significance was calculated based on top 3 peptides. For p-value and fold change calculations, the data were further processed using a custom script, as previously described (*10*).

**Immunoprecipitation assays (co-IP) in live cells.**
For endogenous protein immunoprecipitation using protein A/G beads, cell lysates with equal amounts of protein were diluted with PBS and incubated with protein A/G beads (pre-washed three times with binding buffer, 50 mM Tris, 150 mM NaCl, 0.2% Triton, pH=7.5) for 2 h at 4 ˚C along with the protein-specific antibody at the vendor-suggested dilution. The beads were washed three times with binding buffer (50 mM Tris, 150 mM NaCl, 0.2% Triton, pH=7.5). The enriched proteins were eluted with acidic elution buffer (100 mM glycine, 0.1% Triton, pH 2.8) before neutralizing with 1M Tris (pH 8), according to the manufacturer's instructions.

**AlphaFold-Multimer prediction and analysis.**
We performed in silico screening using AlphaFold-Multimer program on the ColabFold platform as previously described (*11-13*). In brief, AlphaFold-Multimer calculations were performed using AlphaFold-Multimer v3 on ColabFold v1.5.2 (*14*) using NVIDIA A100 GPUs with sequence alignment generated through MMseqs2 and HHsearch. Predictions were generated in a combination of the paired and unpaired multiple sequence alignment, 20 recycles to generate 5 independent unrelaxed models. Sequences were obtained directly from UniProt database. All ranks are examined by both prediction confidence and accuracy. Models with an average predicted local distance difference threshold (pLDDT)>50 and minimum predicted alignment error (PAE)<15 Å were considered.

Predicted AlphaFold-Multimer binary complexes were further scored using predicted DockQ score (pDockQ) and buried solvent accessible surface area (BSASA) (*15, 16*). pDockQ scores were generated to indicate the interface accuracy quantitatively (0 is the worst and 1 is the best) with ≥0.23 cutoff value for direct binary contact as previously described (*15*). BSASA is defined as defined as $\Delta SASA_{AB} = SASA_A + SASA_B - SASA_{AB}$, where a 1.4 Å radii rolling probe was used to calculate solvent accessible surface area for all non-hydrogen, non-monoatomic ion atoms in chains A and B (*14*)(*17, 18*).

**Cell co-culture.**

In order to activate cell-cell recognition, a mixture of two cells were prepared and counted. In the case of BiTE systems, target cell line (HEK293T-Flag-EGFR, HEK293T-Flag-CDCP1, SKBR3) were first plated and let attach to the plate for 8 h. Jurkat NFAT-GFP was then added at a 2.5:1 effector: target ratio. Indicated concentration of BiTE construct was added and incubated for 20 h before the cells were harvested for flow cytometry or on-cell biotinylation assay. In the case of the CAR model, Jurkat-CAR and K562-CD19 were plated at a 2.5:1 effector: target ratio and incubated for 20 h before the cells were harvested for flow cytometry or on-cell biotinylation assay.

**Software.**

Data were analyzed and visualized using Microsoft Excel (v16.22) and GraphPad Prism (v8.0.1), in addition to software listed by each experiment. NMR data were analyzed using MestReNova (v14.0.1). DNA and protein sequences were analyzed using Geneious (v10.0.7). FACS data were analyzed by FlowJo (v10.6.1). Proteomics data were analyzed by PEAKS online (Xpro 1.6) and FragPipe powered by MSFragger (v3.7). Images were made using ImageStudioLite (v5.2.5), Adobe Illustrator (v22.1) and BioRender. Structural assignments to site-specific modifications and structural comparison were performed using ChimeraX (v1.6.1). AlphaFold-Multimer prediction was performed on ColabFold v1.5.2 and visualized using ChimeraX (v1.6.1). Code for BSASA calculation is publicly available: https://github.com/ajipalar/guide_bsasa (commit 7541e6f3ba89a0089b7f01c8792a2f356264cd68).

**Statistical analysis.**

Statistical analyses (unpaired Student's *t*-tests) were performed using GraphPad Prism. Data were derived from at least three biological replicate experiments and presented as the mean ± s.d., P≤ 0.05, **P≤ 0.01, ***P ≤ 0.001, ****P ≤ 0.0001 and n.s., not significant.

#### Synthetic procedures.

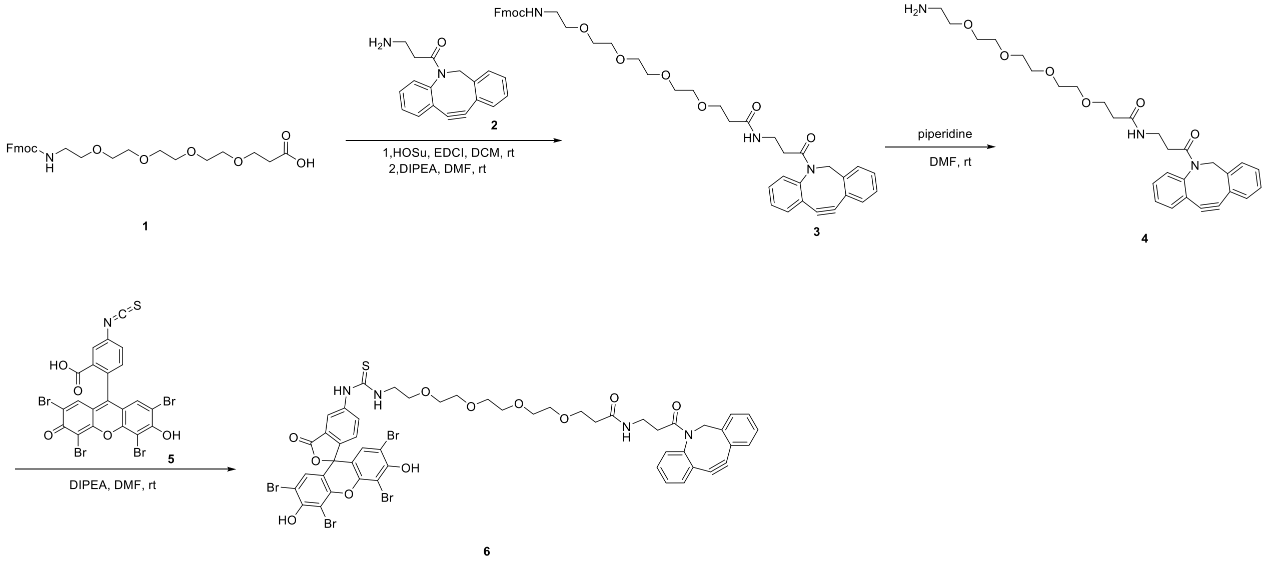


**Scheme 1. Synthesis of DBCO-PEG$_4$-EY (6).**

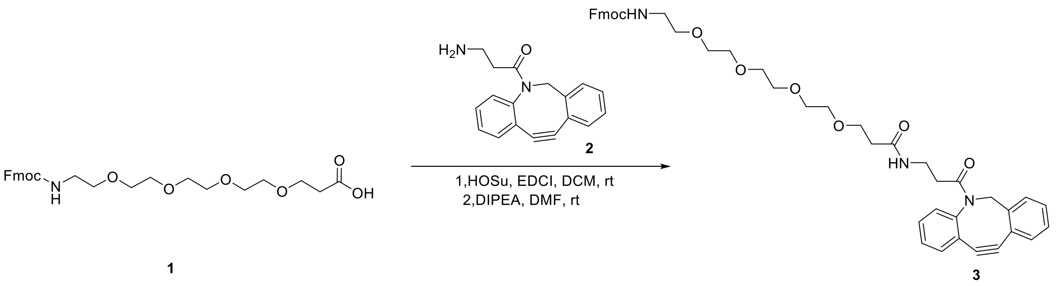

#### Synthesis of compound 3.

To a solution of 1-(9H-fluoren-9-yl)-3-oxo-2,7,10,13,16-pentaoxa-4-azanonadecan-19-oic acid **1** (974 mg, 2 mmol, 1.0 equiv.) in DCM (20 mL) was added HOSu (460 mg, 4 mmol, 2.0 equiv.) and EDCI (764 mg, 4 mmol, 2.0 equiv.) under $N_2$ atmosphere. The mixture was stirred at 25 ˚C for 2 h. The mixture was poured into saturated sodium chloride aqueous solution (26.2 wt%) and extracted with EtOAc. The organic phase was dried over anhydrous magnesium sulfate, filtered, and concentrated in vacuum to afford a residue, which was dissolved in DMF (5 mL). DIPEA (516 mg, 4 mmol, 2.0 equiv.) was then added to the resulting solution, followed by compound **2** (552 mg, 2 mmol, 1.0 equiv.). The mixture was stirred at 25 ˚C for 2 h. The crude reaction mixture was purified by reverse phase HPLC (eluting with 0-65% acetonitrile in water with 0.01% TFA) to give compound **3** as a yellow solid (650 mg, 0.87 mmol, yield: 43%). m/z = 746.1 $[M+H]^+$

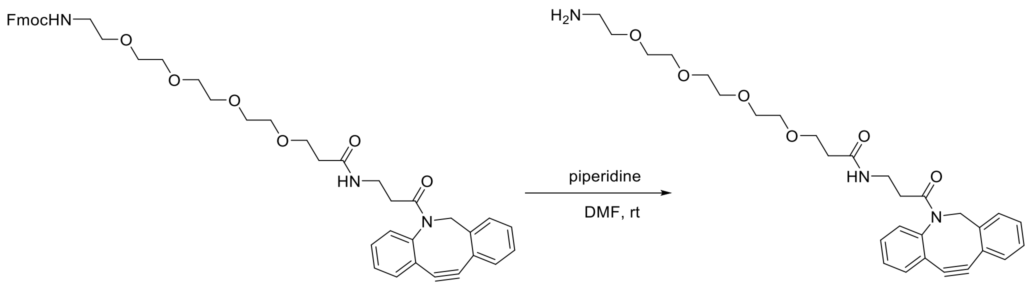


**Synthesis of compound 4.**

To a solution of **3** (650 mg, 0.87 mmol, 1.0 equiv.) in DMF (5 mL) was added piperidine (148 mg, 1.75 mmol, 2.0 equiv.) under $N_2$ atmosphere. The reaction mixture was stirred at 25˚C for 2 h and directly purified by reverse phase HPLC (eluting with 0-65% acetonitrile in water with 0.01% TFA) to give **4** as a yellow solid (400 mg, 0.76 mmol, yield: 88%). ESI m/z = 524.2 $[M+H]^+$

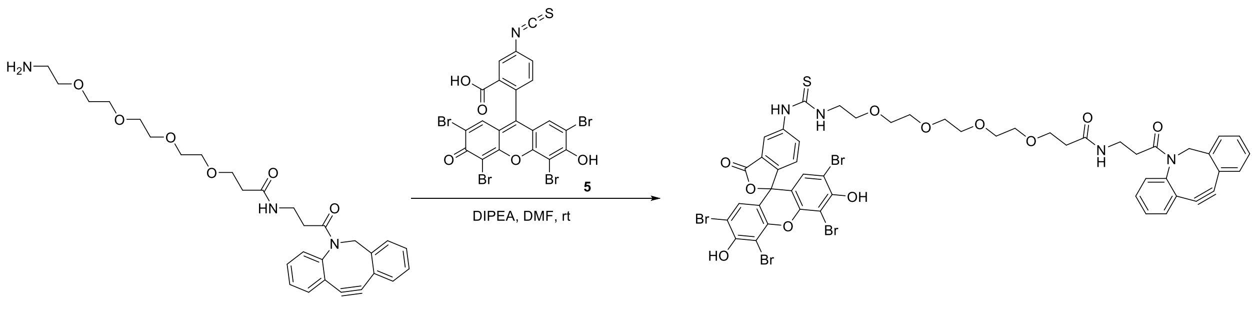

4

6

**Synthesis of compound 6.**

To a solution of **4** (300 mg, 0.56 mmol, 1.0 equiv.) in DMF (3 mL) was added DIPEA (145 mg, 1.12 mmol, 2.0 equiv.) and **5** (400 mg, 0.56 mmol, 1.0 equiv.). The mixture was stirred at rt for 2 h and directly purified by reverse phase HPLC (eluting with 0-65% acetonitrile in water with 0.01% $NH_4HCO_3$) to give 6 as a red solid (200 mg, 0.16 mmol, yield: 30%). Characterization was performed by ChemPartner. m/z = 615.0 $[M/2 +H]^+$

$^1$H NMR (400 MHz, DMSO-$d_6$) δ 10.08 (brs, 1H), 8.25 (s, 1H), 8.18 (s, 1H), 7.88 (d, *J* = 4.0 Hz, 1H), 7.71-7.63 (t, *J* = 5.6 Hz 1H), 7.63-7.58 (m, 2H), 7.50-7.45 (m, 3H), 7.42-7.28 (m, 5H), 7.04 (s, 2H), 5.03 (d, *J* = 14.0 Hz, 1H), 3.72-3.67 (m, 2H), 3.64 (s, 1H), 3.61-3.58 (m, 3H), 3.57-3.52 (m, 4H), 3.50-3.43 (m, 6H), 3.45-3.38 (m, 3H), 3.15-3.17 (m, 1H), 2.98-2.78 (m, 1H), 2.47-2.41 (m, 1H), 2.16 (t, *J* = 6.4 Hz, 2H), 1.84-1.76 (m, 1H)

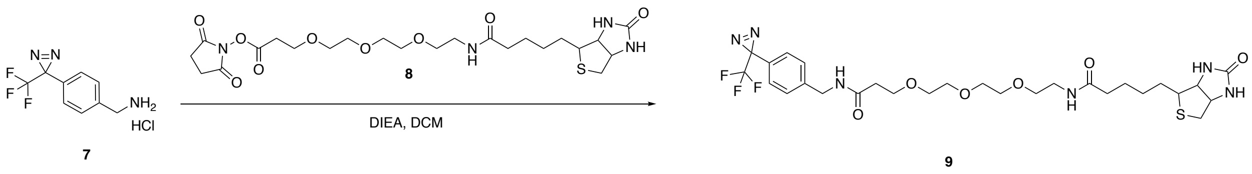


**Scheme 2. Synthesis of diazirine-PEG$_3$-biotin (9)**

To a solution of [4-[3-(trifluoromethyl)diazirin-3-yl]phenyl]methanamine,hydrochloride (20 mg, 0.08 mmol, compound **7**) and biotin-PEG$_3$-NHS ester (47.7 mg, 0.088 mmol, compound **8**) in dichloromethane (1 mL) was added ethylbis(propan-2-yl)amine (31 mg, 0.24 mmol). The reaction mixture was stirred for 1 h at 25 °C. The reaction mixture was concentrated under reduced pressure. The residue was purified by C18 column chromatography eluted with acetonitrile:water (with 0.1% trifluoroacetic acid) (0~40%) to afford diazirine-PEG$_3$-biotin (7.64 mg, 13% yield, compound **9**) as a white solid. Characterization was performed by Medicilon. m/z = 645.2 $[M+H]^+$

$^1$H NMR (400 MHz, DMSO-$d_6$) δ 8.43 - 8.40 (m, 1H), 7.84 - 7.81 (m, 1H), 7.39 - 7.37 (m, 2H), 7.25 - 7.23 (m, 2H), 6.41 - 6.35 (m, 2H), 4.30 - 4.29 (m, 3H), 4.13 - 4.10 (m, 1H), 3.62 (t, *J*=8.0 Hz, 2H), 3.49 (s, 8H), 3.38 (t, *J*=4.0 Hz, 2H), 3.20 - 3.15 (m, 2H), 3.11 - 3.08 (m, 1H), 2.59 - 2.56 (m, 1H), 2.38 (t, *J*=4.0 Hz, 2H), 2.06 (t, *J*=4.0 Hz, 2H), 1.63 - 1.56 (m, 2H), 1.51 - 1.40 (m, 3H), 1.33 - 1.26 (m, 2H)

**NMR spectrum for DBCO-PEG$_4$-EY (6).**

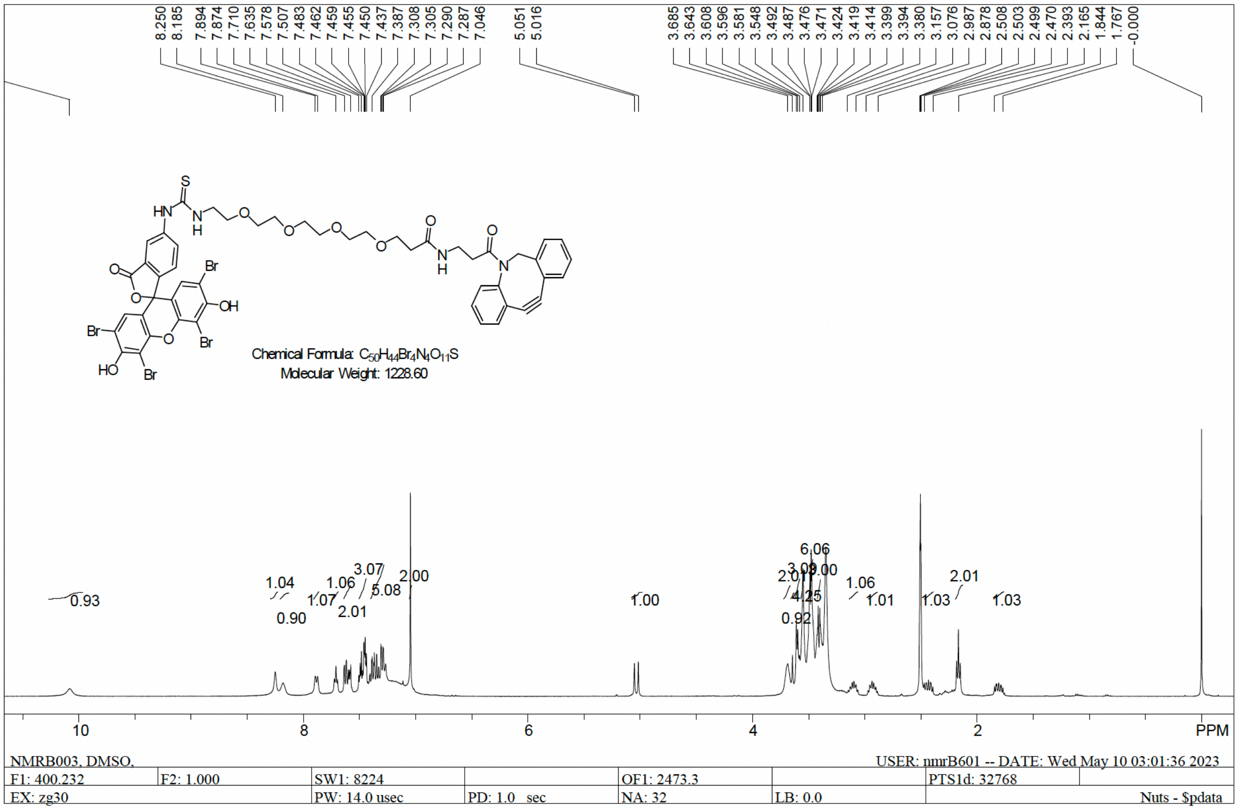

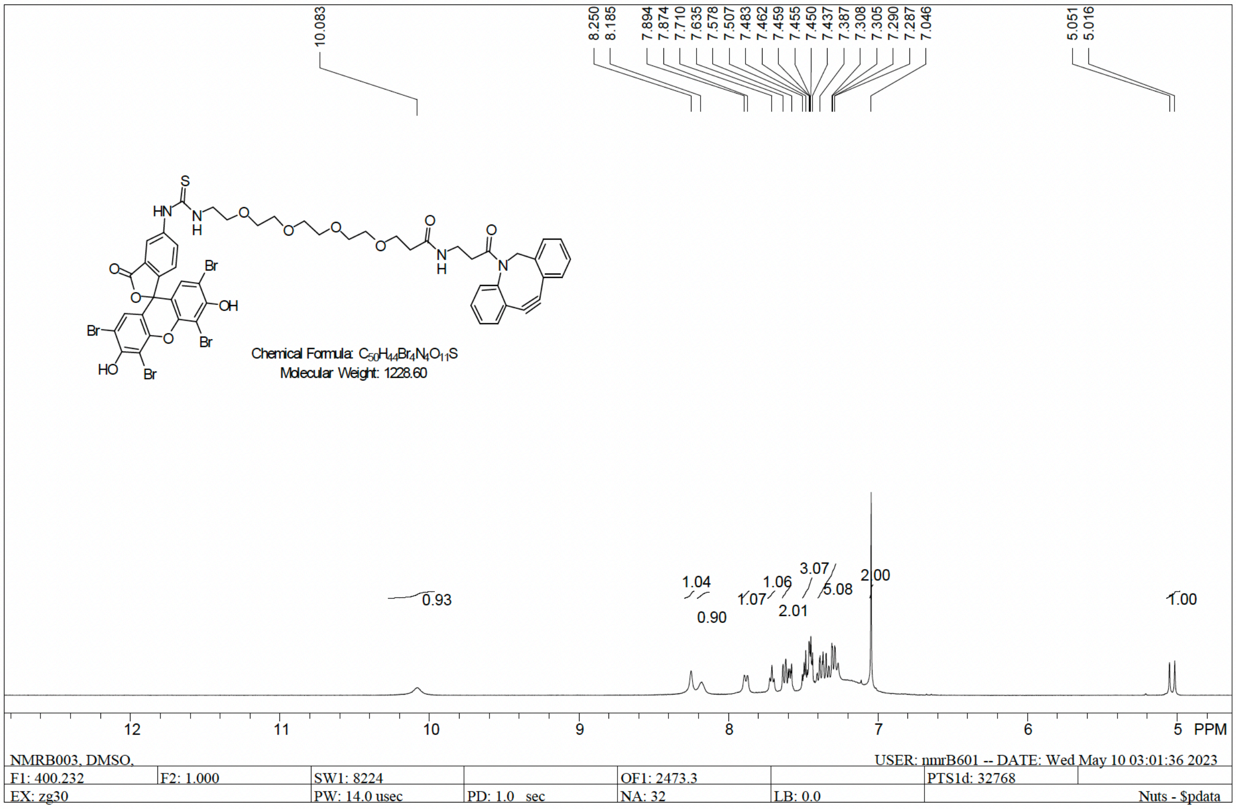

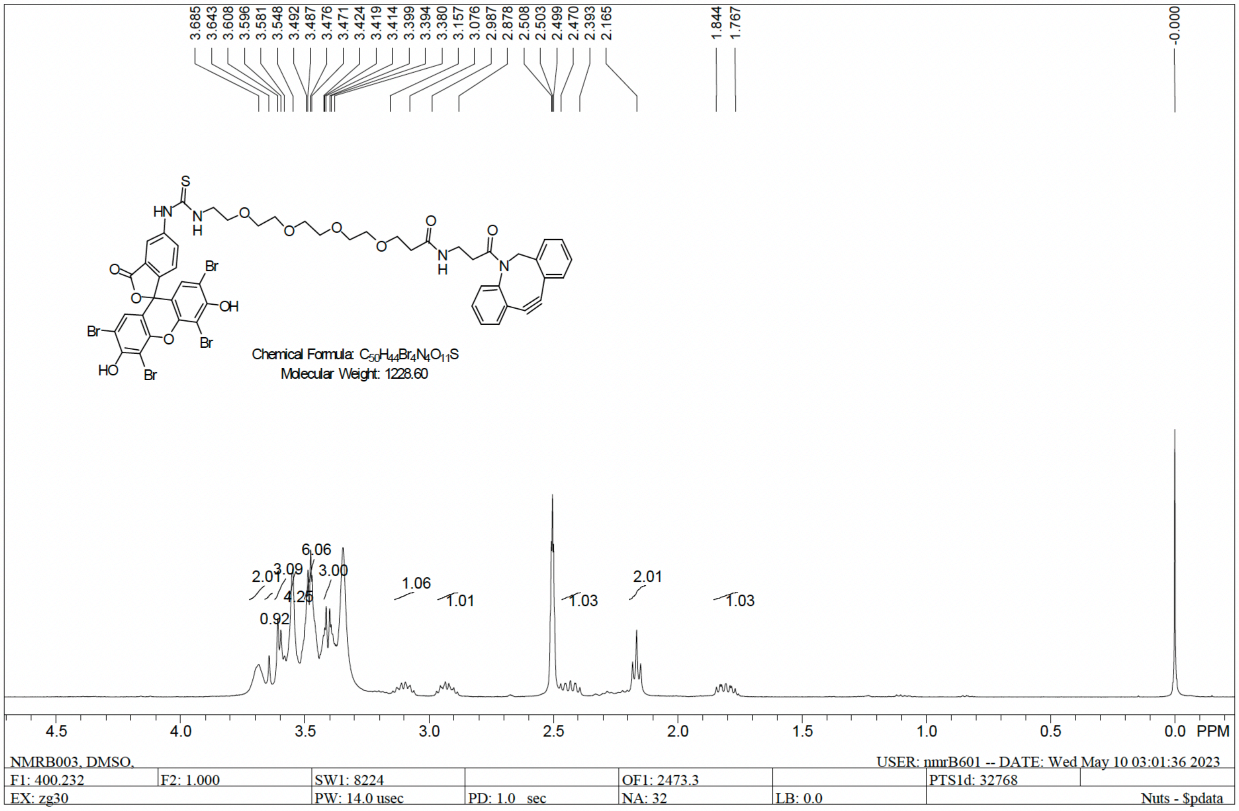
3.685 3.643 3.608 3.596 3.581 3.548 3.492 3.487 3.476 3.471 3.424 3.419 3.414 3.399 3.394 3.380 3.157 3.076 2.987 2.878 2.508 2.503 2.499 2.470 2.393 2.165 1.844 1.767 -0.000
S
HN
N
H
O
O
O
O
O
N
H
O
N
Br
OH
Br
O
Br
HO
Br
O
Chemical Formula: C50H44Br4N4O11S
Molecular Weight: 1228.60
2.01 3.09 6.06 3.00 4.25 0.92 1.06 1.01 1.03 2.01 1.03
4.5 4.0 3.5 3.0 2.5 2.0 1.5 1.0 0.5 0.0 PPM
NMRB003, DMSO,
USER: nmrB601 -- DATE: Wed May 10 03:01:36 2023
F1: 400.232
F2: 1.000
SW1: 8224
OF1: 2473.3
PTS1d: 32768
EX: zg30
PW: 14.0 usec
PD: 1.0 sec
NA: 32
LB: 0.0
Nuts - $pdata

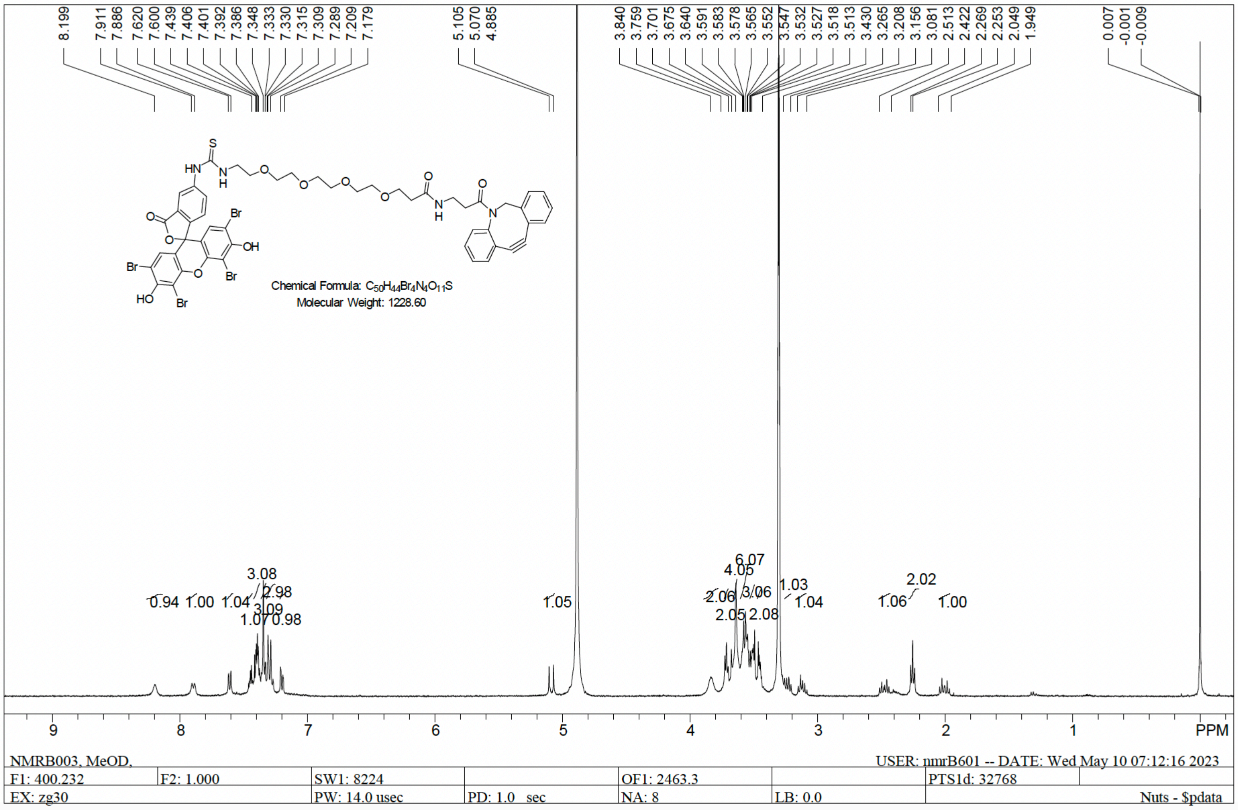
8.199 7.911 7.886 7.620 7.600 7.439 7.406 7.401 7.392 7.386 7.348 7.333 7.330 7.315 7.309 7.289 7.209 7.179 5.105 5.070 4.885 3.840 3.759 3.701 3.675 3.640 3.591 3.583 3.578 3.565 3.552 3.547 3.532 3.527 3.518 3.513 3.430 3.265 3.208 3.156 3.081 2.513 2.422 2.269 2.253 2.049 1.949 0.007 -0.001 -0.009
S
HN
N
H
O
O
O
O
O
N
H
O
N
Br
OH
Br
O
Br
HO
Br
O
Chemical Formula: C50H44Br4N4O11S
Molecular Weight: 1228.60
0.94 1.00 1.04 3.08 2.98 3.09 1.07 0.98 1.05 2.06 4.05 6.07 3.06 2.05 2.08 1.03 1.04 1.06 2.02 1.00
9 8 7 6 5 4 3 2 1 PPM
NMRB003, MeOD,
USER: nmrB601 -- DATE: Wed May 10 07:12:16 2023
F1: 400.232
F2: 1.000
SW1: 8224
OF1: 2463.3
PTS1d: 32768
EX: zg30
PW: 14.0 usec
PD: 1.0 sec
NA: 8
LB: 0.0
Nuts - $pdata

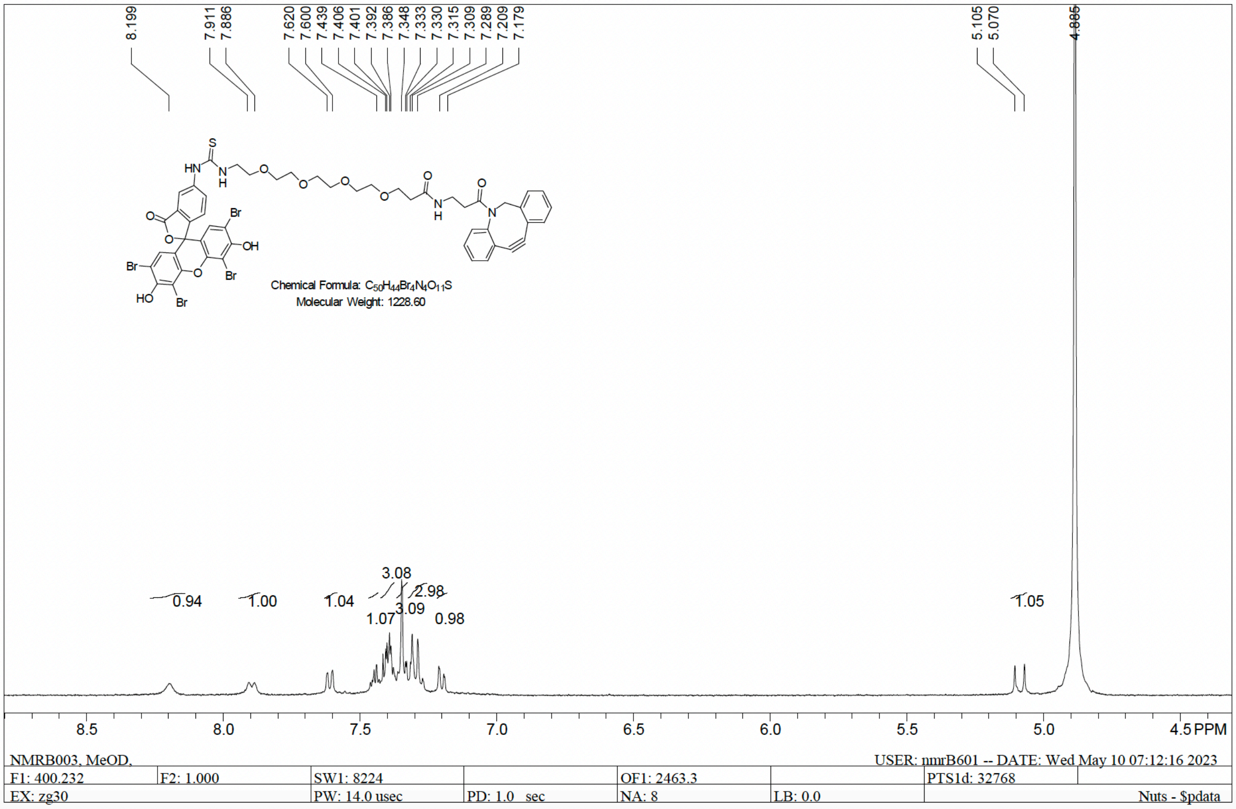
Chemical Formula: $C_{50}H_{44}Br_4N_4O_{11}S$
Molecular Weight: 1228.60
NMRB003, MeOD,
USER: nmrB601 -- DATE: Wed May 10 07:12:16 2023
F1: 400.232
F2: 1.000
SW1: 8224
OF1: 2463.3
PTS1d: 32768
EX: zg30
PW: 14.0 usec
PD: 1.0 sec
NA: 8
LB: 0.0
Nuts - $pdata
PPM

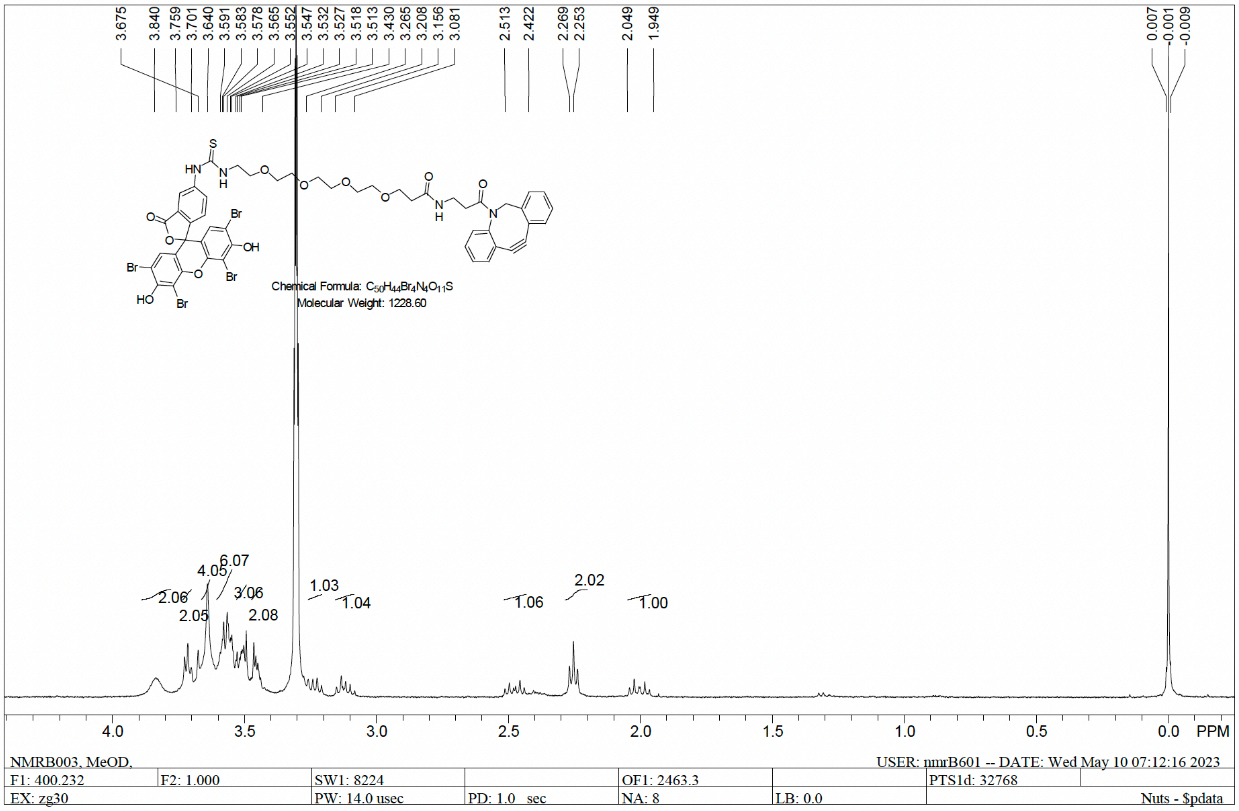
Chemical Formula: $C_{50}H_{44}Br_4N_4O_{11}S$
Molecular Weight: 1228.60
NMRB003, MeOD,
USER: nmrB601 -- DATE: Wed May 10 07:12:16 2023
F1: 400.232
F2: 1.000
SW1: 8224
OF1: 2463.3
PTS1d: 32768
EX: zg30
PW: 14.0 usec
PD: 1.0 sec
NA: 8
LB: 0.0
Nuts - $pdata
PPM

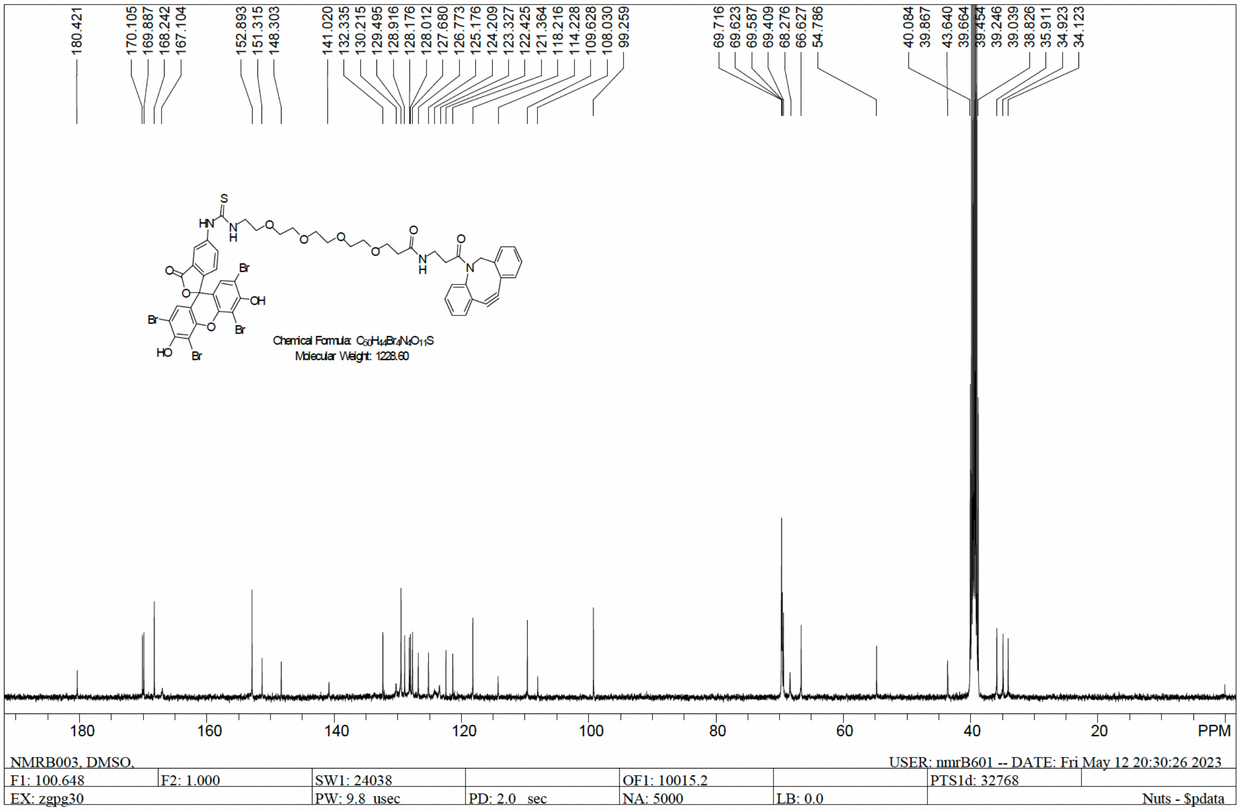
180.421
170.105
169.887
168.242
167.104
152.893
151.315
148.303
141.020
132.335
130.215
129.495
128.916
128.176
128.012
127.680
126.773
125.176
124.209
123.327
122.425
121.364
118.216
114.228
109.628
108.030
99.259
69.716
69.623
69.587
69.409
68.276
66.627
54.786
40.084
39.867
43.640
39.664
39.454
39.246
39.039
38.826
35.911
34.923
34.123
Molecular Weight: 1228.60
PPM
NMRB003, DMSO,
USER: nmrB601 -- DATE: Fri May 12 20:30:26 2023
F1: 100.648
F2: 1.000
SW1: 24038
OF1: 10015.2
PTS1d: 32768
EX: zgpg30
PW: 9.8 usec
PD: 2.0 sec
NA: 5000
LB: 0.0
Nuts - $pdata

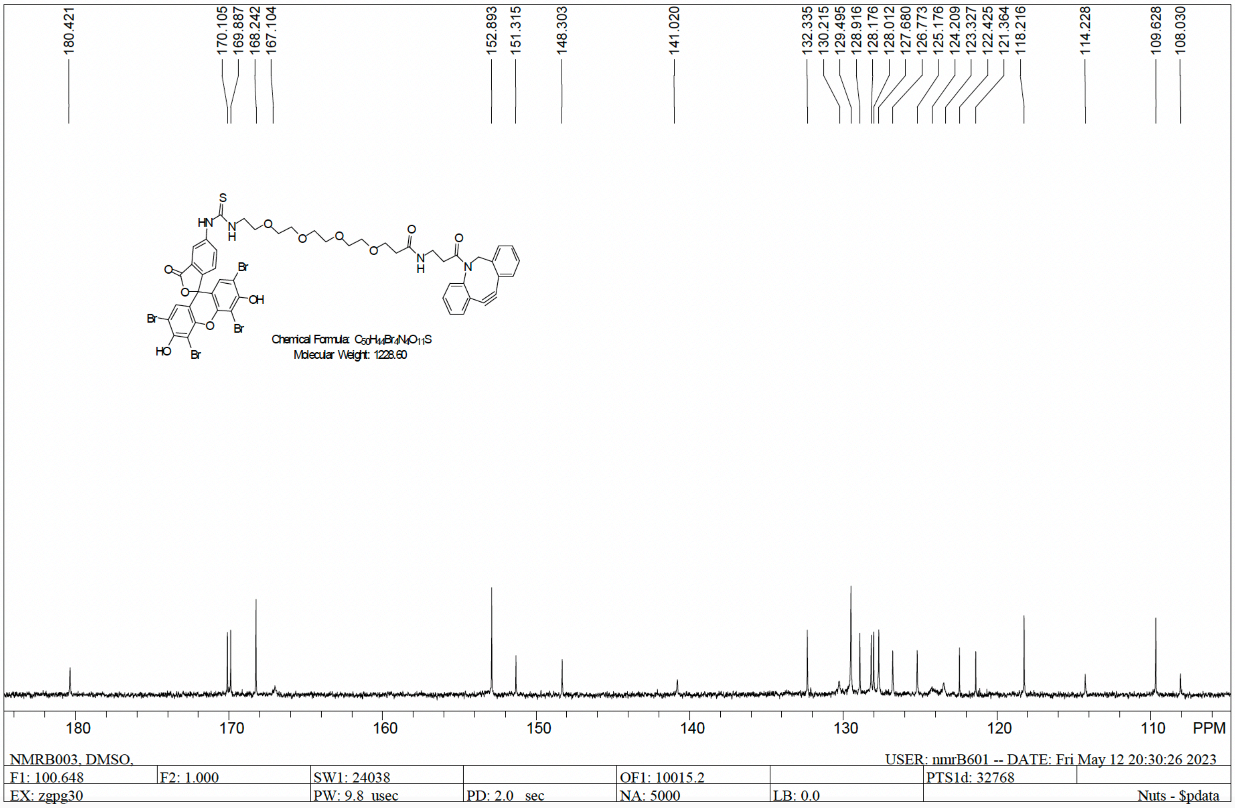
180.421
170.105
169.887
168.242
167.104
152.893
151.315
148.303
141.020
132.335
130.215
129.495
128.916
128.176
128.012
127.680
126.773
125.176
124.209
123.327
122.425
121.364
118.216
114.228
109.628
108.030
Molecular Weight: 1228.60
PPM
NMRB003, DMSO,
USER: nmrB601 -- DATE: Fri May 12 20:30:26 2023
F1: 100.648
F2: 1.000
SW1: 24038
OF1: 10015.2
PTS1d: 32768
EX: zgpg30
PW: 9.8 usec
PD: 2.0 sec
NA: 5000
LB: 0.0
Nuts - $pdata

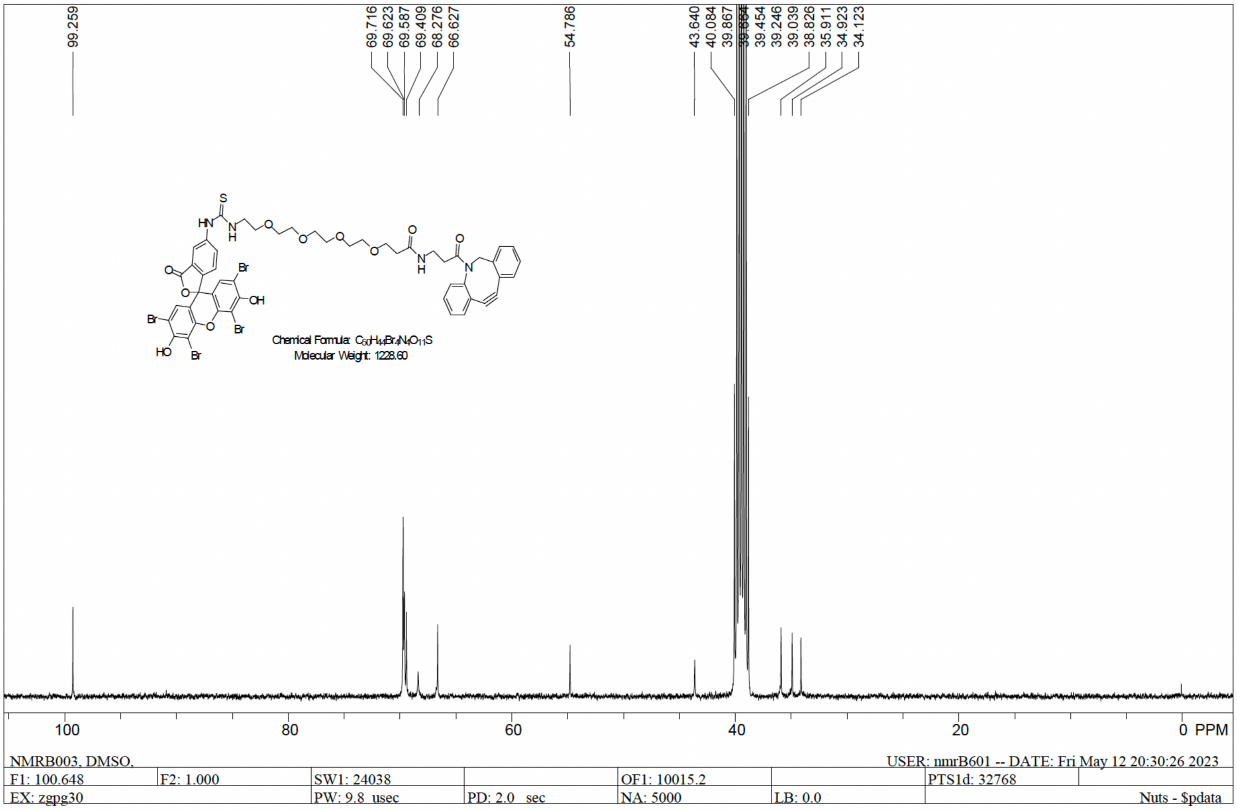

99.259
69.716
69.623
69.587
69.409
68.276
66.627
54.786
43.640
40.084
39.867
39.454
39.246
39.039
38.826
35.911
34.923
34.123
Chemical Formula: C50H40Br4N4O11S
Molecular Weight: 1228.60
100
80
60
40
20
0 PPM
NMRB003, DMSO,
USER: nmrB601 -- DATE: Fri May 12 20:30:26 2023
F1: 100.648
F2: 1.000
SW1: 24038
OF1: 10015.2
PTS1d: 32768
EX: zgpg30
PW: 9.8 usec
PD: 2.0 sec
NA: 5000
LB: 0.0
Nuts - $pdata
